# On the adaptive behavior of head-fixed flies navigating in two-dimensional, visual virtual reality

**DOI:** 10.1101/462028

**Authors:** Hannah Haberkern, Melanie A. Basnak, Biafra Ahanonu, David Schauder, Jeremy D. Cohen, Mark Bolstad, Christopher Bruns, Vivek Jayaraman

## Abstract

A navigating animal’s sensory experience is shaped not just by its surroundings, but by its movements within them, which in turn are influenced by its past experiences. Studying the intertwined roles of sensation, experience and directed action in navigation has been made easier by the development of virtual reality (VR) environments for head-fixed animals, which allow for quantitative measurements of behavior in well-controlled sensory conditions. VR has long featured in studies of *Drosophila melanogaster*, but these experiments have typically relied on one-dimensional (1D) VR, effectively allowing the fly to change only its heading in a visual scene, and not its position. Here we explore how flies navigate in a two-dimensional (2D) visual VR environment that more closely resembles their experience during free behavior. We show that flies’ interaction with landmarks in 2D environments cannot be automatically derived from their behavior in simpler 1D environments. Using a novel paradigm, we then demonstrate that flies in 2D VR adapt their behavior in a visual environment in response to optogenetically delivered appetitive and aversive stimuli. Much like free-walking flies after encounters with food, head-fixed flies respond to optogenetic activation of sugar-sensing neurons by initiating a local search behavior. Finally, by pairing optogenetic activation of heat-sensing cells to the flies’ presence near visual landmarks of specific shapes, we elicit selective learned avoidance of landmarks associated with aversive “virtual heat”. These head-fixed paradigms set the stage for an interrogation of fly brain circuitry underlying flexible navigation in complex visual environments.

## Introduction

Animals in their natural habitat often navigate complex visual environments to forage for food, find mates and escape inhospitable conditions or predators. Insects, in particular, are among the animal kingdom’s most skilled navigators, and have been the focus of decades of field and laboratory studies [1-6]. These studies have produced important insights into the range of navigational algorithms that insects use in different naturalistic contexts [7-9]. However, it is difficult to closely monitor and flexibly control an animal’s sensory experience in natural settings; a shortcoming that is addressed by the complementary approach of studying behavior in virtual reality (VR) [10]. VR approaches enable the creation of environments with customized rules for how the virtual sensory surroundings change in response to an animal’s actions, and have found wide application in neuroscience across species [11-19].

In *Drosophila*, VR paradigms have been used to study the behavior of flying [11, 20, 21] and walking [22-24] flies. However, most previous studies of navigational behaviors in tethered flies have been limited to 1D environments, in which the fly only controls its orientation relative to a fixed circular visual panorama, and its translational movements are disregarded. The conclusions drawn from the fly’s behavior in such reduced environments can be challenging to translate to more realistic settings in which a fly’s location —and not just its heading— matters. Indeed, most studies of freely moving flies highlight the 2D nature of their behavior, whether in flight [25, 26] or walking [27, 28]. Inferring movement trajectories through 2D space is challenging in tethered flying flies, but this problem is easier to solve in a tethered walking preparation [29, 30].

Here we explore the navigational strategies of head-fixed flies in a versatile 2D VR system. We exploit the flexibility of VR to explore how head-fixed flies change their interaction with visual landmarks depending on the visual stimulus conditions and past experiences. The latter makes use of optogenetics to activate sensory receptors, giving flies an experience of taste and heat in an otherwise purely visual VR. While optogenetic activation of the appropriate sensory receptors does not fully substitute for genuine consumption of sugar or actual heat, this strategy has experimental advantages. It allows for experiments in which an animal’s response to such sensory experiences can be studied without affecting its physical state —level of satiation and body condition. Optogenetic activation of sensory neurons has previously been used to study olfactory processing [31, 32], CO_2_ avoidance [33] and thermotaxis [34]. Direct activation of sensory and dopaminergic neurons has also been used to replace reward or punishment in odor conditioning assays [35-38]. Further, in a recent report, freely moving flies were shown to avoid areas in which their bitter taste receptors were optogenetically activated, although it was not clear if flies could acquire a conditioned place preference as a result of such experience [39]. Inspired by such studies we developed behavioral paradigms to study adaptive, visually-guided navigation in moderately complex 2D VR environments. We show that encounters with optogenetically-induced sweet taste trigger exploratory behavior similar to those seen in freely walking flies upon encounters with real food or optogenetic substitutes [27, 40-42]. Motivated by the demonstration that freely walking flies can learn to use visual cues to navigate to a cool spot in an aversively hot 2D environment [43], we also trained head-fixed flies to avoid visual landmarks that were associated with optogenetic activation of the heat-sensing pathway. These VR paradigms for head-fixed flies clear a path towards understanding the neural underpinnings of exploratory and learned navigation in flies.

## Results

### A 2D visual VR system for head-fixed walking flies

To explore 2D navigation in tethered flies, we built a VR system that combined an existing spherical treadmill for head-fixed walking flies with a projector-based panoramic visual display (**Fig. 1A**, **Fig S1A-C**, Methods). As in past 1D VR experiments [23, 44], we glued flies to a thin wire tether and suspended them above an air-supported ball such that the fly’s walking maneuvers resulted in rotations of the ball (**Fig. 1A** inset, **B, C**). These ball rotations were tracked at high temporal resolution by optical mouse sensor chips [22]. To translate the fly’s measured walking movements into simulated movement through a 2D virtual visual environment, we developed a C++ program (“FlyoVeR”, based on a program developed for generating a VR environment for rodents [45], Methods, **Fig. S1A**). 2D environments were designed with 3D-modeling software and loaded into FlyoVeR (Methods). The VR system permits the fly to walk towards and around a landmark in these environments (**Fig. 1D**, Methods), with the landmark increasing in size as the fly approaches it (**Fig. 1F**). As landmarks, we used simple, salient, rotationally symmetric, geometric shapes such as cones and cylinders (**Fig. 1F** and **Fig. S1G,I**). We limited the number of collisions with the impenetrable landmarks by using virtual scenes with sparsely distributed landmarks and avoided confining the animal inside virtual walls, as our purely visual VR cannot adequately model collisions. To increase the fly’s sampling of the visual landmarks, we arranged them in large periodic grids, so that the fly repeatedly encountered identical visual scenes. We will refer to such a world made from cone-shaped landmarks as a *single-landmark forest* (**Fig. 1H, Fig. S1E,H**, Methods). The radial symmetry of the virtual scene permitted a compact, “landmark-centric” description of the fly’s position based on the fly’s distance and heading relative to a nearby landmark (**Fig. 1E,G**). For the analysis of walking trajectories, we exploited periodicity and sparse placement of landmarks to generate “collapsed” trajectories, pooling points in the VR that corresponded to the same visual environment (**Fig. 1I**). To visually separate the landmarks, we used virtual fog, hiding any objects beyond a user-defined distance from the fly (**Fig. 1J, Fig. S1G,I**).

**Figure 1:**
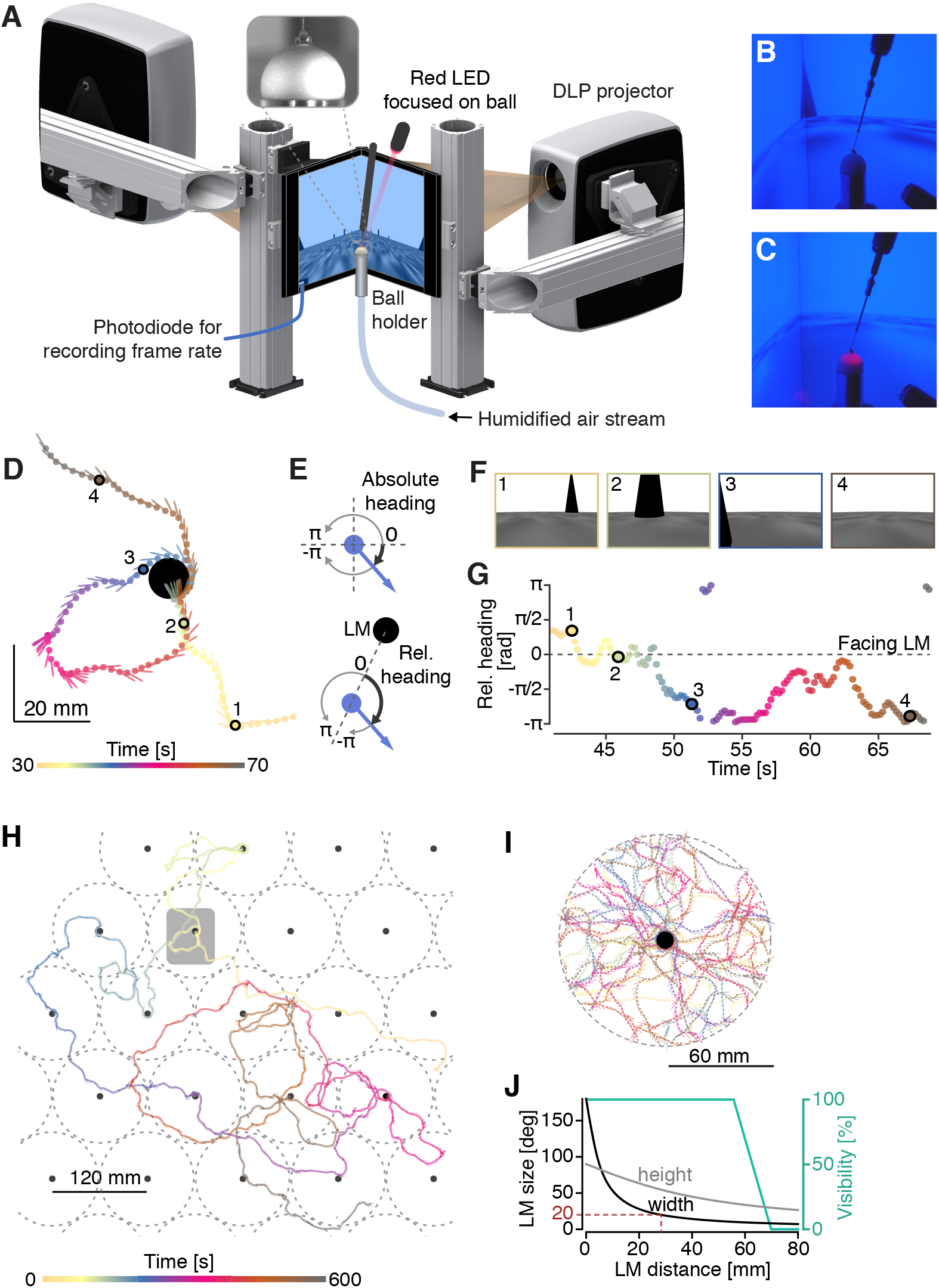
A 2D virtual reality (VR) system for head-fixed walking fruit flies. (**A**) Schematic of the projector-based visual display and its position relative to the tethered walking fly. The inset shows an image taken from behind the fly with an IR camera used for calibration and fly positioning. (**B, C**) Photographs of a fly in VR viewed from the side. The IR LEDs used for illuminating the ball are visible in the lower right corner. In (C), the fly receives optogenetic stimulation (red light). (**D**) Example trace of a fly exploring a cone (black circle) in a virtual 2D plane. The position fly (center of mass) is indicated with a dot and a short line marks the fly’s viewing direction. The progression of time is color-coded. (**E**) Illustration of absolute and relative heading angles. (**F**) Rendered images of the scene from the point of view of the fly at the four time points marked by black circles in the trace fragment in (D). Note that these rendered images are for illustration and do not contain the perspective corrections applied to images that were projected onto the screen during the experiment. See Fig. S2 b-d for screenshots of projected images. (**G**) Relative heading angle of the fly with respect to the landmark in the trace fragment shown in (D). A relative heading angle of 0 corresponds to facing the landmark. Note that the gap in the panoramic screen behind the fly means that the landmark is only visible for relative heading angles smaller than ⅔ π or larger than -⅔ π. Color-code as in (D). (**H**) Trajectory of a fly exploring a periodic world with cone-shaped landmarks (*single landmark forest*) over the course of a 10 min trial. Shaded box: section of the trajectory shown in (D). Grey dashed circles: circular area around each landmark that was included in the analysis. (**I**) Trajectory from (H) after “collapsing” the trajectory fragments within a 60 mm radius circle around each landmark to one circular reference “arena” (60 mm radius) with a centrally placed landmark. (**J**) Schematic illustrating how a landmark’s (10 mm wide, 40 mm high cone) angular dimensions change as a function of distance, as seen from the fly’s point of view. The blue line indicates visibility of the landmark through the virtual fog (100%: full visibility, 0%: zero visibility). The distance at which the cone has an angular width of 20 is highlighted. DLP, digital light processing; LM, landmark.

### Validation of closed-loop feedback: stripe fixation

As a first step toward validating our VR system, we asked whether the feedback loop between the fly’s movements and the visual stimulus was sufficiently fast and the displayed landmarks sufficiently salient for tethered flies to perform a well-established naïve visual behavior — stripe fixation [46, 47]. In this closed-loop assay, the fly only controls its angular orientation relative to a circular panorama with a single stripe (inset **Fig. 2A**), while its distance relative to the panorama is fixed. Under this condition, tethered walking flies actively keep the stripe in their frontal field of view (FOV), particularly at high temperatures [44]. We found that, as expected, over the course of a 10-minute trial, some flies fixated a black stripe in their frontal FOV resulting in a single peak around 0 in the relative heading distribution (frontal fixation, **Fig. 2A**). Across genotypes, females consistently showed higher rates of fixation compared to males (**Fig. 2B, C**, **Fig. S2 C-F**, see Methods for details) and fixating flies, preferentially kept the stripe in their frontal FOV (**Fig. 2B**, see Methods for details). The precision of frontal fixation varied across genotypes (**Fig. 2B**, distribution wider for WTB hybrid flies compared to DL flies), potentially reflecting a difference in walking behavior: DL flies displayed lower translational speeds, and those WTB hybrid flies that walked with similar speeds showed clearer frontal fixation (**Fig. S2A,B**). These experiments confirmed that our VR system permits stripe tracking behavior, and revealed systematic behavioral differences between genotypes and sexes.

**Figure 2:**
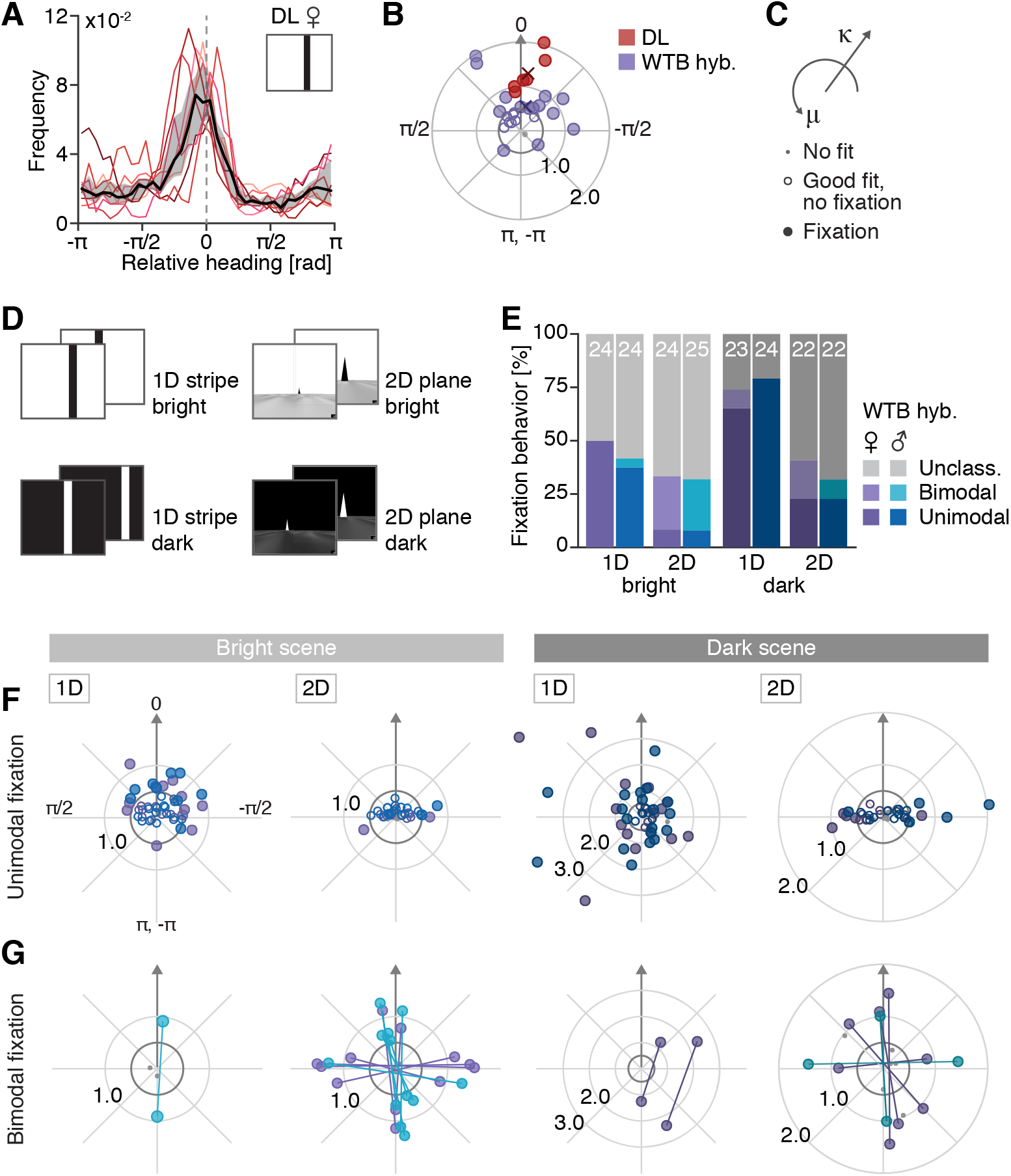
Fixation behavior is affected by world dimension and contrast polarity. (**A**) Relative heading distributions of individual female DL flies (n = 6) that fixated a black stripe in a bright scene are shown as colored lines. The black line and grey shaded region indicate median and interquar-tile range (IQR, between the 25th and 75th quartile). Relative heading angle bin size: 10 °. (**B**) Polar “fixation plot” (legend in C) showing the angular location of the fixation peak and the fixation strength based on von Mises fits for female DL (n = 8, red) and female WTB hybrid flies (n = 22, purple). The grey arrow points toward the frontal position in the fly’s field of view. Only flies that were walking sufficiently during the trial were included: 14 out of 22 WTB hybrid flies and 6 out of 8 DL flies (details in Methods). **(C)** Guide to the polar “Fixation plots” in (B,F,G), which quantify fixation behavior based on von Mises function fits. The location parameter (μ) is plotted on the circumferential axis and the shape parameter (κ) on the radial axis. Markers (dot, empty or filled circle) indicate categorization of trial based on fit. (**D**-**G**) Fixation behavior of WTB hybrid flies (male and female, n = 18 for each) across four types of virtual worlds. (**D**) Illustration of the four scenes: 1D stripe in bright scene, 2D cones in bright scene, 1D stripe in dark scene and 2D cones in dark scene. (**E**) Frequency of bimodal and unimodal fixation. For each bar, the white number indicates the number of flies that were walking and hence included into the analysis (25 flies measured in each case). (**F,G**) Fixation plots visualizing fixation directions based on a von Mises fit in the bright (left) and dark (right) scenes. Data from male and female flies are color-coded as in (E). (**F**) Fixation directions for unimodal fixation. (**G**) Fixation directions for bimodal fixation. The two fitted location parameters (μ1, μ2) corresponding to the same measurement are connected by a line. Only data from flies that walked for at least 20% of a trial was included in the analyses in this figure. Note that in experiments with DL, the wings were glued, while in experiments with WTB hybrid flies, the wings were cut. DL, Dickinson lab strain; WTB hybrid, hybrid generated from WTB and empty-Gal4 line.

### Fixation behavior is affected by world dimensionality and contrast polarity

When an animal explores a 2D environment, it can not only control its orientation relative to visual landmarks, as in a 1D VR world, but it can also move towards and away from them. To investigate how a simple visual orientation behavior such as stripe fixation translates to 2D environments, we tested fixation behavior of WTB hybrid flies in 1D (stripe VR) and 2D (single-landmark-forest VR) environments with at most one salient landmark. We also examined the role of contrast polarity —whether landmarks appear as dark objects on a bright background (*dark-on-bright*), as in most natural scenes, or as bright objects on a dark background (*bright-on-dark*), as is frequently used in fly VR experiments.

Testing fixation across four scenes varying in world dimensionality and contrast polarity (**Fig 2D**; see Methods), we found that rates of fixation varied widely (**Fig 2E**), even though walking rates were similar. Fixation was more frequent in *bright-on-dark* conditions and in 1D compared to 2D scenes (**Fig. 2E**). In 2D trials relative heading angle distributions often showed two fixation peaks, a behavior which we termed “bimodal fixation” (**Fig. 2E, Fig. S2E-F**, Methods). Intuitively, bimodal fixation may be expected in a 2D world, as it is consistent with periods of approach (landmark in the front) and departure (landmark in rear, **Fig. S2E-H**). Moreover, fixation as measured here may be less prominent in 2D than 1D environments because movement toward a landmark during translation tends to push the landmark out of the frontal FOV unless it is perfectly centered. Besides the rate of fixation, the distribution of preferred fixation angles also varied with scene type: While most flies showed frontal fixation of a black stripe (**Fig. 2F** far left), fixation of the bright stripe occurred across the entire FOV (**Fig. 2F** center right). Curiously, flies that showed unimodal fixation of landmarks in 2D scenes kept the landmark in their lateral FOV, which corresponds to a circling trajectory around the landmark (**Fig. 2F** center left and far right**, Fig. S2G,I**), but would correspond to maintaining a fixed heading if the landmarks were far away. In flies that showed bimodal fixation, the two fixation peaks were typically located at opposite locations within the fly’s FOV (π offset, **Fig. 2G**). Frequently, the two fixation peaks were either in the front or back, corresponding to straight trajectories from landmark to landmark (**Fig. S2G,H**), or at the side, corresponding to counterclockwise and clockwise circling around the landmarks in this environment. Results obtained with other genotypes, DL and WTB, were similar (**Fig. S3B-F**). These features of fixation behavior also persisted across a range of temperatures and were not affected by manipulations aimed at rendering flies flightless (**Fig. S3G-I**). Thus, consistent with previous findings in tethered flight [48, 49], scene contrast polarity and scene brightness affect fixation behavior both quantitatively and qualitatively.

### Flies interact with landmarks in VR much as they do during free behavior

In the real world, collisions with objects result in mechanosensory feedback, which is completely absent when flies encounter virtual objects in our purely visual VR. To assess the impact of this limitation of our VR system, we compared the behavior of flies navigating in 2D VR to that of freely walking flies. We built a large free walking arena (**Fig. S4A-C**) where flies could interact with a real landmark under lighting conditions similar to those in VR. Importantly, and in contrast to previous studies [50], we prevented flies from climbing on the landmark (Methods). In free walking experiments (**Fig. 3A-C**), we let individual flies explore the arena with a single black cone-shaped landmark placed in the center (**Fig. S4A**). We then compared their walking trajectories to walking traces in VR from tethered flies exploring the *single-landmark forest* world (**Fig 3D-F**). Walking velocities were similar, though translational velocities were higher in freely walking flies and variability across flies higher in VR (**Fig S4D-I**). Several flies showed multiple landmark approaches and departures over in both real world and VR (**Fig. 3A,D**), matched by a general preference for frontal fixations of the landmark (**Fig. 3C,F**, **Fig. S4F**). Importantly, under both conditions flies showed an increased residence near the landmark (**Fig. 3B,C,E,F**). This was not a consequence of landmarks physically blocking the fly’s path —or, in VR, caused in VR by landmark impenetrability. Flies in conditions identical to the *single-landmark forest* but with all landmarks invisible, yet impenetrable, did not show the previously described elevated residency around the landmarks (**Fig. 3G**). This could not be explained simply by the flies being less active (**Fig. 3H**) or by altered walking velocities in the absence of visible landmarks (data not shown). Rather, flies made more visits to the visible landmarks compared to the invisible landmarks (**Fig. 3I**) and stayed there for longer (data not shown), indicating that flies actively steered toward visible landmarks.

**Figure 3:**
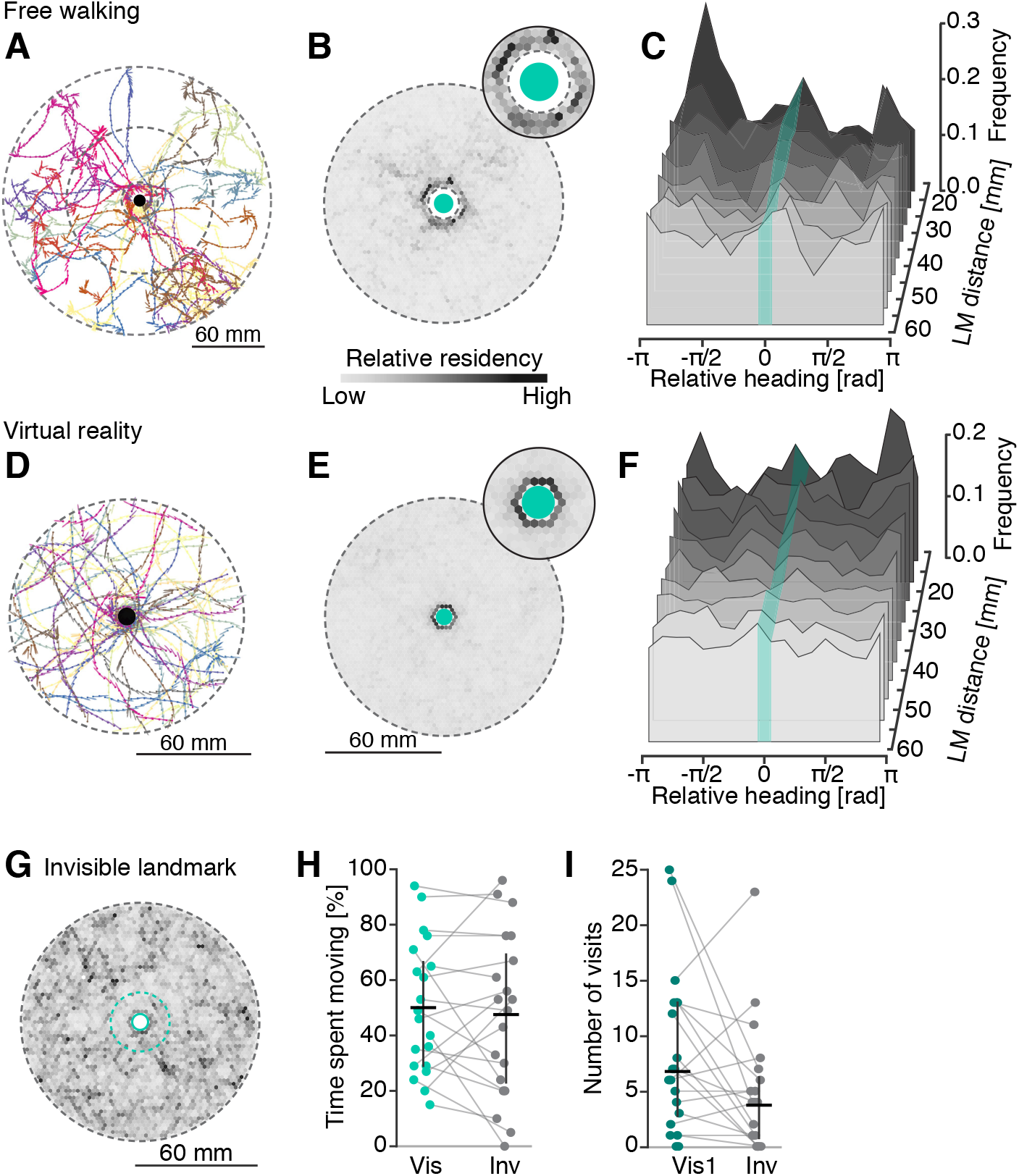
Landmark interaction in freely walking and head-fixed flies. (**A**-**F**) Comparison of landmark interaction of n = 20 freely walking flies in an arena with a single central physical landmark (**A-C**) versus n = 20 flies exploring the *single landmark forest* scene VR (**D-F**). For the analysis of free walking data, only trajectory fragments within 15 mm and 60 mm radial distance from landmark (area between the inner two dashed circles in (A)) were included. (**A**, **D**) Walking trajectory of a single fly over a 10 min trial visualized as in Fig. 1I. (**B**, **E**) Residency histogram of collapsed trajectories in Cartesian coordinates for pooled data from free walking (B) and VR (E, data additionally pooled over three 10 min trials) experiments. Only time points when a fly was moving were considered. The count was normalized for each histogram and color-coded with darker shades indicating high residency. Turquoise circle: landmark position. Insets: zoom in on area around landmark. (**C**, **F**) Visualization of residency in landmark-centric polar coordinates for free walking (C) and VR (F, three 10 min trials pooled) data. Data was binned according to landmark distance and for each 10 mm wide bin the relative heading distribution was computed (angle bin size: 20 °). The resulting distributions for each bin are visualized as staggered planes with varying grey shading as a qualitative indicator for the landmark distance. Turquoise stripes: angular position, but not the width, of the landmark. (**G**-**I**) Interaction with visible and invisible virtual landmarks in VR (n = 20). (**G**) Residency histogram in Cartesian coordinates for trials with invisible landmarks. Dashed turquoise circle: visit radius (15 mm) used for subsequent analysis. (**H**) Percentage of trial time flies spent moving in trials with visible (average across the three trials) and invisible landmarks. Data from single flies shown as dots with a grey line connecting corresponding measurements. The mean and IQR are shown in black. (**I**) Total number of landmark visits in first trial with visible and in trial with invisible landmarks. All data from female WTB hybrid flies. LM, landmark.

### Using optogenetics to study context-dependent navigation

Flies modify their behavior depending on context, whether that is an encounter with food [27, 40] or changes in environmental conditions, such as temperature [6, 51]. Introducing context into a VR environment presents challenges —food can be difficult to present to flies without leaving traces on the surface that the fly walks on, and exposure to persistently high temperature can damage flies physically. We thus sought to add context to the otherwise purely visual virtual environment by using optogenetic activation of appropriate sensory pathways [40].

### Flies initiate local search behavior upon transiently tasting “virtual sugar”

When flies encounter food they initiate a local search behavior [27, 40-42] that has been suggested to rely on path integration [27]. To examine the as yet unexplored role of visual cues in this local search behavior, we sought to evoke the behavior in 2D VR using transient (200 ms) optogenetic stimulation of taste receptors. Hungry flies are known to slow down and extend their proboscis upon encounters with sweet-tasting food [52]. This behavior likely depends, in part, on the Gr64f gustatory receptor, which is expressed in many sweet-sensing neurons [53-55]. We first verified that transiently activating these neurons with optogenetics in populations of flies exploring a free walking arena (see Methods) induced them to slow down almost immediately (**Fig. S5**). Transient activation of neurons expressing Gr64f (“virtual sugar” stimulus) in single flies in VR produced similar reductions in walking speed (**Fig. 4D,E**, Methods).

**Figure 4:**
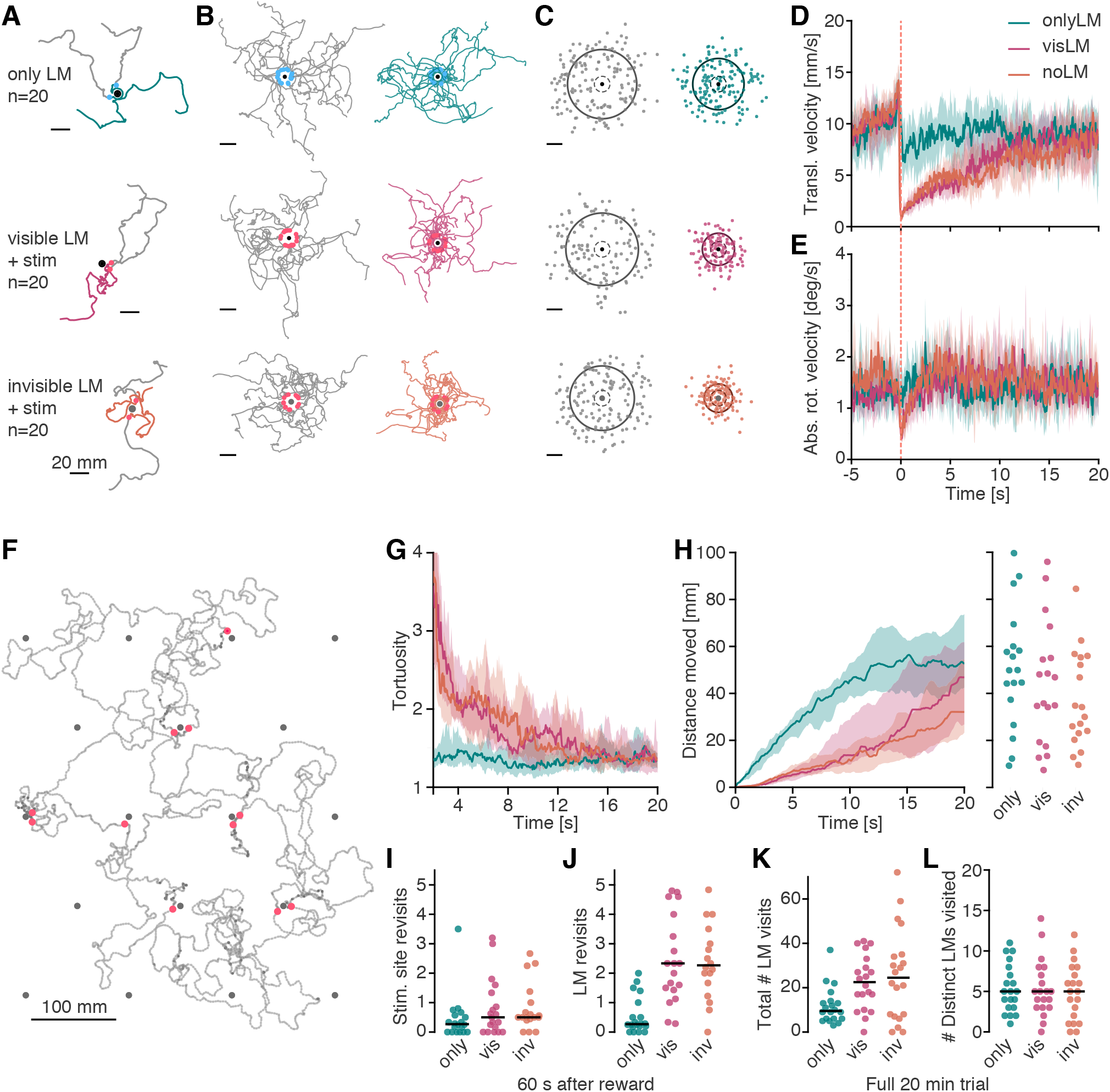
Local search behavior in VR after optogenetic activation of sugar sensing neurons. Local search behavior was investigated in starved (24h) female Gr64f > ChrimsonR flies in three VR conditions: visible landmarks only (single-object forest, data color coded in teal throughout the figure), optogenetic stimulation around the visible landmarks (magenta), and optogenetic stimulation around invisible landmarks (orange). (**A**) Paths of 150 mm length before (grey) and after (colored) optogenetic stimulation. Two trajectories from the same fly are shown for each condition. Sample sizes indicated next to the panel refer to the full group size (n = 20 each). The position of the landmark (visible or invisible) is marked by a black dot. (**B**) Paths of 150 mm length before (grey, left) and after (colored, right) optogenetic stimulation from all flies (if available). For each fly, the trajectory around the first optogenetic stimulus encounter is shown. Colored dots mark the point of the light stimulus on the left panels. (**C**) Center of mass of all 150 mm paths (including those shown in B) before (left) and after (right). Grey circles: mean distance of center of mass. Dashed circles: visit radius (10 mm). (**D, E**) Average change in translational and absolute rotational (E) velocity triggered on the optogenetic stimulation. (**F**) Example trajectory of a fly in the VR with optogenetic stimulation, but no visible landmarks (full 20 min). Dark grey dots: invisible landmark locations. Orange dots: stimulation events. (**G**) Tortuosity measured as the path length over the radial distance moved. Data at each time point corresponds to tortuosity measurement in the preceding 2 s window. (**H**) Radial distance moved away from the optogenetic stimulation site. Left: average across flies over time. Right: median distance moved after 20 s for each fly. (**I,J**) Stimulation site (I) and landmark (J) revisits in a 60 s window following optogenetic stimulation. Each dot represents the mean number of revisits (10 mm radius) per fly. Median across flies shown by black bar. (**K,L**) Total number of landmark visits (K) and number of distinct landmarks that were visited (L) over the course of the full 20 min trial. Each dot represents the count per fly, the black bar shows the median across flies. Thick lines in (D,E,G,H) mark the median across flies and the shaded region marks the interquartile range. Data was first averaged (median) across trajectory fragments per fly and subsequently across flies.

We then tested the effect of pairing the *virtual sugar* stimulus with landmarks in VR. We hypothesized that after experiencing *virtual sugar* flies would increase their exploration of the space around landmarks. Further, we thought that if flies showed local search in VR, its accuracy could be increased in the presence of landmark cues. We tested this in starved flies exploring the *single-landmark forest* world. If landmarks were visible, but no *virtual sugar* was provided, flies approached landmarks, but also quickly departed from them (**Fig. 4A-C** top row), as seen before. By contrast, if flies transiently tasted virtual sugar when reaching a visible landmark, they not only slowed down, but also showed a significant increase in the tortuosity of their walking trajectories (**Fig. 4A,C** middle row, **G**), switching to a local search behavior resembling that observed in recent studies in freely walking flies [27, 40]. This combination of slowdowns and increased turning also resulted in flies spending more time near landmarks (**Fig. 4C** middle, **H**). However, the same adaptations in walking behavior were observed upon *virtual sugar* stimulation when no visible landmark cues were provided (**Fig. 4A,B** bottom row, **G,H**), suggesting that landmark cues were dispensable for this behavior. Indeed, flies returned to the source of the initial *virtual taste* in both visual and non-visual conditions (**Fig. 4I,J**). Note that returns to the local (invisible or visible) landmark within the forest of landmarks were more frequent than returns to the initial source, potentially due to flies encountering the impenetrable landmarks during the search phase and then finding their paths impeded. Pairing visual landmarks with *virtual sugar* stimulation resulted in a larger number of landmark visits across the forest than the presentation of visual landmarks alone (**Fig. 4K**). All three experimental groups of flies sampled the virtual space to a similar degree over the course of the full 20 min trial, as measured by the total number of different landmarks visited (**Fig. 4F,L**). Thus, flies responded with transient changes in behavior upon activation of sugar sensing neurons, initiating bouts of local exploration following *virtual sugar* sensation, a behavior that did not require visual landmarks.

### Flies avoid “virtual heat” when walking freely and in VR

Following the observation that the presence of landmarks did not markedly improve the accuracy of local search behavior in VR, we next sought to explore the role of visual landmarks in avoidance of an aversive stimulus. In particular, we asked whether flies could learn to avoid salient visual landmarks that had been associated with an aversive stimulus. The extensive history of fly visual learning experiments with heat and electric shocks, and the fact that flies have exquisite temperature sensitivity and show robust avoidance of high environmental temperatures [56], led us to explore the effects of pairing an aversive heat stimulus with our visual VR. We generated *“virtual heat”* stimuli by optogenetically activating heat sensing neurons targeted by the hot cell-Gal4 (HC-Gal4) line [57]. We used the free walking quadrant assay to screen optogenetic stimulation intensities (**Fig. S6A-D**) and quantified the induced avoidance response as the fraction of flies residing in the stimulated quadrants (**Fig. 5B,C**). We compared *virtual heat* avoidance in response to three stimulation levels in male and female flies (**Fig. 5C**). Flies robustly avoided even low intensities of stimulation, which likely only activated the peripherally located HC neurons in the fly’s antenna and not the central neurons captured in the HC-Gal4 expression pattern (**Fig. 5A**). At low stimulation intensity, we did also not detect avoidance responses in control flies (**Fig. 5C,** non-retinal control in light grey), suggesting that the red light was not in itself aversive. We concluded that optogenetic activation of HC neurons could be used as an effective aversive stimulus in walking flies. Because virtual heat avoidance appeared to be more pronounced in male flies, we performed all VR experiments in male HC-Gal4 > ChrimsonR flies.

**Figure 5:**
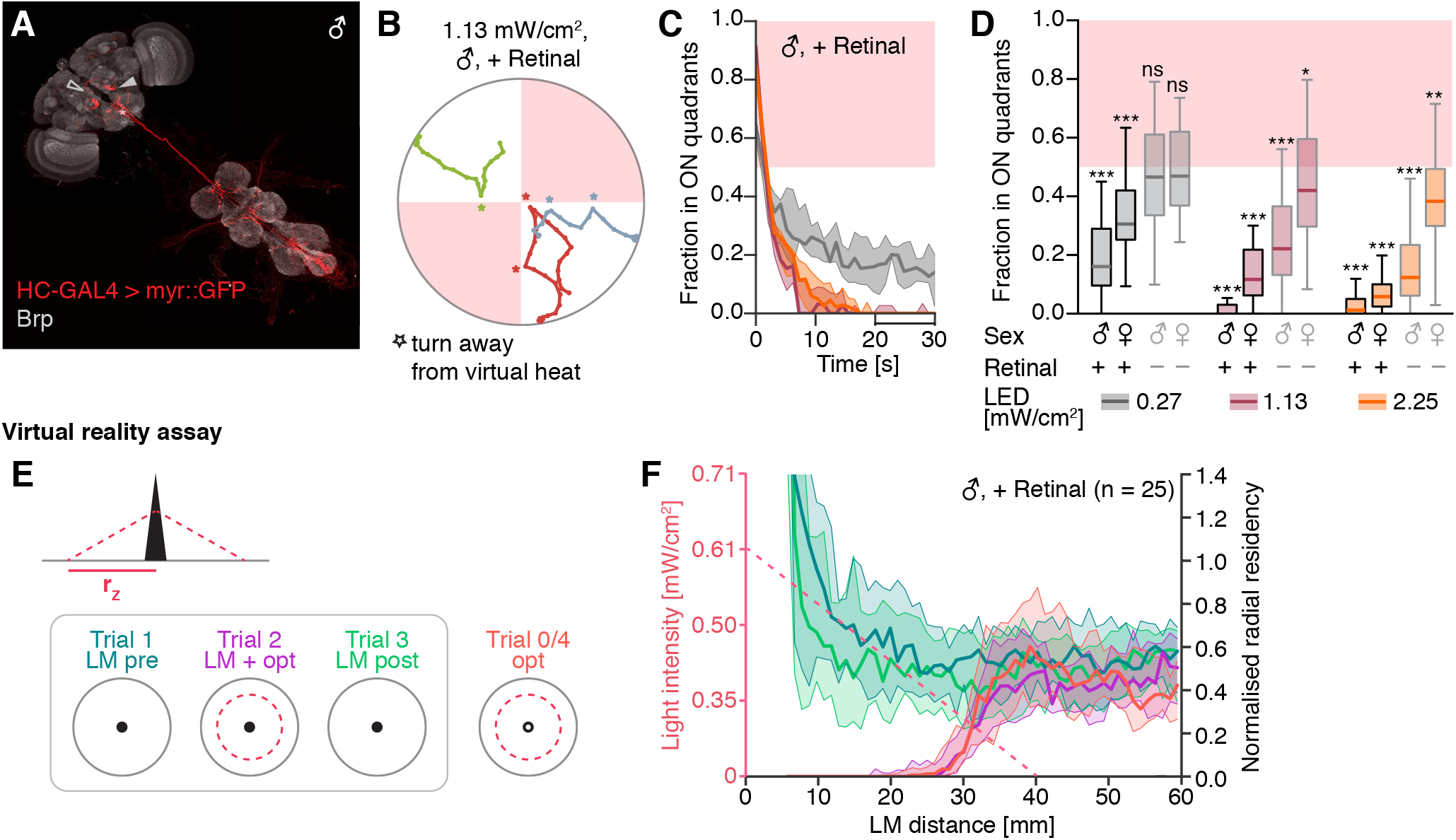
Avoidance of optogenetically induced “virtual heat” in freely walking flies. (**A**) Expression pattern of Hot cell (HC)-GAL4 in the central nervous system. The cell bodies of HC neurons are located in the antennae and have been removed during the dissection, but the dendritic arborizations are visible in the proximal antennal protocerebrum (PAP, filled arrowhead). Additionally, HC-GAL4 labels putative ascending neurons (asterisk). In some brains, a neuron arborizing in the fan-shaped body was also labelled (empty arrowhead). Background staining with nc82 antibody against Bruchpilot (Brp). Panel shows a montage of two separate images of the brain and the VNC. (**B**) Short fragments of walking traces from three male HC-GAL4 > ChrimsonR flies in the free walking optogenetic quadrant assay (fly identity color-coded). Asterisks mark turns away from the illuminated quadrants (red-shaded regions). (**C**) Fraction of male HC-GAL4 > ChrimsonR flies residing in the two illuminated quadrants over the course of the 30 s long stimulation block. The median (thick line) and IQR (shaded region) are shown for three experimental groups with different stimulation light intensities. Light intensities are color-coded as indicated in the legend of (D). Values below 0.5 indicate avoidance of the optogenetic stimulus. (**D**) Comparison of different stimulation protocols in the quadrant assay. For each group the average fraction of flies in the illuminated quadrants measured in the last 10 s of the stimulation repeat (bar in time axis in C) is shown. Statistical significance levels refer to difference from 0.5 (two-sided one-sample t-test). (**E,F**) *Virtual heat* avoidance in VR. (**E**) Illustration of the experimental paradigm for testing landmark-assisted *virtual heat* avoidance in VR. Left: *virtual heat* zone. Right: Trial structure. (**F**) Comparison of the normalized radial residency of male, retinal-fed flies across the four trials illustrated in (E, right). The radial residency is computed as the count of time points within a given radial distance range (relative to the landmark) normalized by the area of that radial distance range. Only time points when the fly was moving faster than 2 mm/s were considered. The dashed salmon-colored line indicates the stimulation light intensity at a given radial distance. Note that the optogenetic stimulus light intensity level is depicted as a continuously varying, but the resolution of the intensity control was limited to steps of 1 % driver current. Significance codes: “ns” p > 0.1, “*” p ≤ 0.05, “**” p ≤ 0.01, “***” p ≤ 0.001.

Next, we generated *virtual heat* zones in VR (**Fig. 5E** left), using stimulation intensities of less than 0.61 mW/cm^2^. We used *virtual heat* zones centered around either visible or invisible landmarks with linearly increasing stimulation intensities, which provided flies with additional information about the shape of the zone. We tested each fly in four trials in the *single-landmark forest* VR (**Fig. 5E** right): two trials with only the visible landmarks, separated by a trial in which visible landmarks were paired with *virtual heat* zones, and a fourth trial where *virtual heat* zones were paired with invisible landmarks. Control flies did not show an avoidance response in VR (**Fig. S6E** center), suggesting that HC activation was indeed responsible for the aversive behavior. To check if flies used the landmarks to avoid *virtual heat* zones in VR, we compared *virtual heat* avoidance between trials with and without visible landmarks by quantifying residency along the radial distance from the closest landmark (**Fig. 5F**, purple and orange profiles). The residency profile around the landmark was similar between the two conditions, indicating that landmark cues were not necessary for this type of avoidance behavior.

Notably, experiencing *virtual he*at zones paired with visible landmarks did not reduce visit rates or residency around landmarks in the absence of *virtual heat* (light and dark green profiles **in Fig. 5F**; **Fig. S6E**), potentially because the flies’ innate drive to approach landmarks overrode any learned aversion.

### Flies distinguish between landmark shapes in VR

We reasoned that any aversive visual conditioning paradigm in VR would need to counter a fly’s strong drive to approach landmarks. Therefore, we devised a paradigm that would shift the fly’s relative preference for different landmark shapes rather than altering its behavior in the presence of a single landmark type. As a prerequisite for such a paradigm, we first established that flies could distinguish between two landmark shapes: a cylinder and a cone. To test flies’ naïve preferences for these landmarks, we created a second type of periodic virtual world, a *“two-landmark forest”* (**Fig. S1H,I, Fig. 6A, Movie 1**). Indeed, when left to explore the *two-landmark forest*, most male WTB hybrid flies showed increased residency around cylinders and higher visit rates to cylinders than cones (**Fig. 6B-E**). Thus, flies were able to distinguish the shapes and selectively approached one type of landmark more frequently.

**Figure 6:**
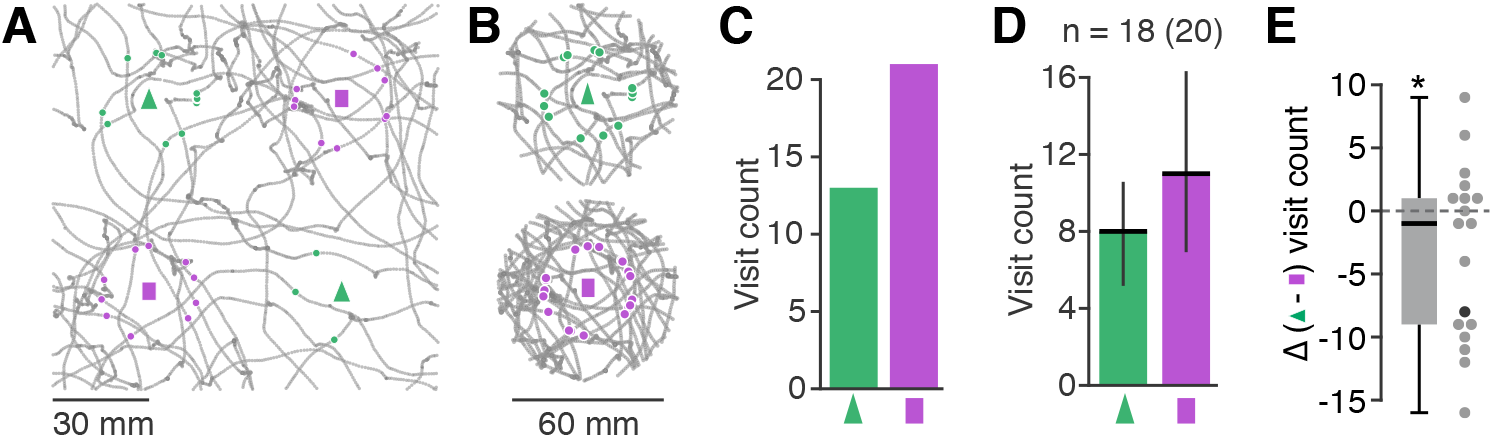
Naïve preference landmark preference in VR. Relative preference of male WTB hybrid flies for cylindrical over conic landmarks in a two-landmark forest VR. **(A)** Example trajectory (collapsed) of a single fly over the 10 min trial. Landmark positions are marked by the green triangle (cone) and magenta rectangle (cylinder). Green and magenta dots mark when the fly entered a radius of 15 mm around a cone or cylinder, respectively (‘visits’). **(B)** Trajectory of the example in (A) further collapsed to area within 30 mm around the two landmark types. Visits illustrated as in (A). **(C)** Resulting visit count for cylinders and cones for the same example fly. **(D, E)** Quantification of visit counts and landmark preference across flies (n = 20 flies measured, n = 18 flies selected based on minimum number of 5 visits to any landmark). **(D)** Average visit count (median and interquartile range). **(E)** Relative preference measured as the difference in the total number of visits to cones minus cylinders (one sample t-test with null hypothesis of mean = 0: p = 0.06423). Black dot marks preference of fly shown in A-C.

### Flies in VR alter their preference for landmark shapes associated with “virtual heat”

Having validated the different components required for a visual conditioning assay, we now paired the *two-landmark forest* (**Fig. S1H,I**) with a *virtual heat* landscape that changed over the course of three consecutive trials (**Fig. 7A**). In all three trials, flies were exposed to a baseline of constant low *virtual heat* that kept flies moving through the environment enough to result in multiple visits to both landmarks. Previous experiments in the *single-landmark forest* environment had shown that a baseline of low levels of *virtual heat* reduced but did not abolish avoidance of *virtual heat* zones (**Fig. S6E** right). In a pre-trial phase, we let flies explore the visual environment and measured the fly’s innate preferences between the two landmarks (**Fig. 7B,C** left). On average, HC > ChrimsonR flies had a naïve preference for cylinders over cones (**Fig. 7C,D** left) as seen previously in WTB hybrid flies. In training trials, *virtual heat* was increased within a small zone around cylinders (“*hot zone*”) and decreased around cones (“*cool zone*”, **Fig. 7A** center, **Movie 2**). In this anti-cylinder training protocol, flies avoided entering the *hot zone*, resulting in a low rate of visits to cylinders, whereas they often stopped at the edge of the *cool zone* near the cone, resulting in an increased residency 10-15 mm away from the cones (**Fig. 7B-C,** center). In the post-trial phase, *cool* and *hot zones* were removed, but on average flies kept visiting cones more frequently than cylinders (**Fig. 7B-D**, right). Thus, pairing a *virtual heat* landscape with the two visual landmarks over the course of 20 minutes of training was sufficient to significantly and consistently alter naïve landmark preferences, quantified as the difference in cone and cylinder visits (**Fig. 7E, Fig. S7D**). This learning effect did not depend on the visit radius chosen (**Fig. S7A-C**). The shift in landmark preference was driven by a combination of increased visit rates to cones and decreased visit rates to cylinders (**Fig. 7E**, compare visit counts). We also tested the reverse training protocol (anti-cone), in which cylinders were paired with *cool zones* and cones with *hot zones* (**Fig. 7F**). In this paradigm flies showed a naïve preference for cylinders and avoided the *hot zones*, but while the time course of the landmark preference mirrored that of the anti-cylinder protocol, the shift in landmark preference before and after training was not significant (**Fig. S7E**). To further exclude the possibility that the observed shift in landmark preference after anti-cylinder training had been induced by mere exposure to *virtual heat*, we designed a control paradigm in which the cool and hot zones were shifted such that they were no longer associated with cone and cylinder positions, respectively (**Fig. 7G**). In this paradigm we no longer observed any shifts in landmark preference (**Fig. 7G, Fig. S7F**). We therefore conclude that the shift in landmark preference observed in our anti-cylinder conditioning paradigm relies on an association of aversive and hospitable *virtual heat* stimuli with the relevant visual landmarks.

**Figure 7:**
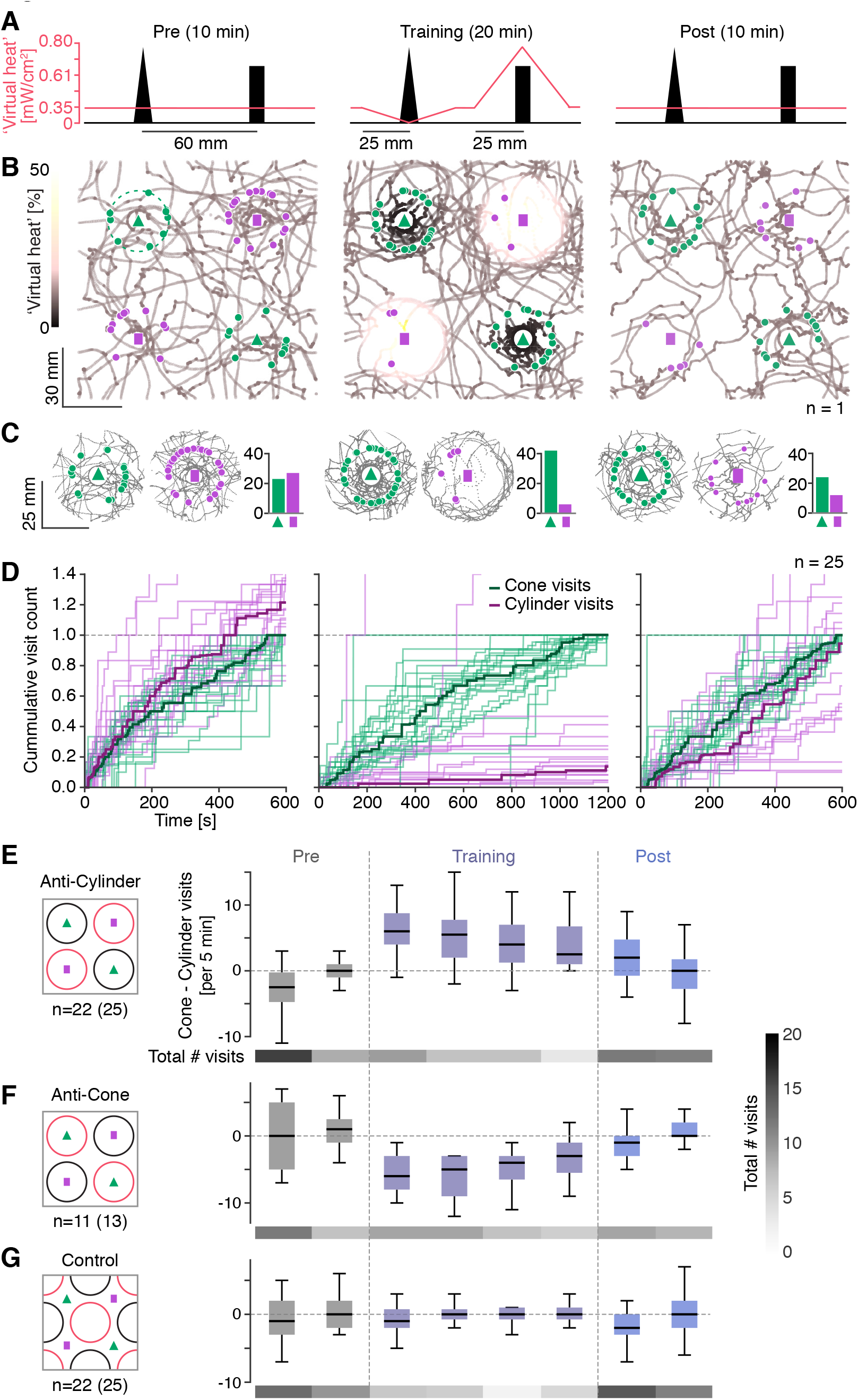
Conditioning of landmark preferences with “virtual heat”. **(A)** Schematic illustrating the anti-cylinder conditioning paradigm consisting of a pre-(left), training-(center) and post-trial (right). A section through the virtual plane cutting along the center of one cone and one cylinder is shown. The salmon line indicates the *virtual heat* level along this section for each trial. (**B**) Collapsed walking trace of a single male HC > ChrimsonR fly across the three trials. The positions of cones are marked by green triangles and those of cylinders by pink rectangles. Each “visit” to a landmark is marked by a pink or green dot for a cylinder or cone visit, respectively. The walking trajectory is color-coded according to the level of *virtual heat* that the fly experienced at the respective location. (**C**) Illustration of the visit preference for a single fly across the three trials (same fly as in B). Collapsed trajectories (grey) within a 25 mm radius around the two landmark types are shown. Overlaid green and pink circles indicate visits to the cone and cylinder, respectively. The bar graph to the right of each subpanel shows the total visit count for the two landmarks (green: cone, pink: cylinder). (**D**) Cumulated visit count over the length of each trial. For each fly, the cumulated count for both cylinder and cone visits is normalized by the maximum number of cone visits. Thin lines: visit counts for individual flies. Thick lines: median. (**E-G**) Comparison of shift in landmark preference across three conditioning paradigms: anti-cylinder conditioning (E, same data as in A-D), anti-cone conditioning (F) and a shifted zone control, in which *virtual heat* zones are not paired with specific landmarks (G). Landmark preference was quantified for each fly in intervals of 5 min as the number of cone visits minus the number of cylinder visits. Data from all flies that made at least 5 visits to any landmark in each trial is presented as boxplots (black line: median, box spans the 25th to the 75th quartile). Sample sizes before and after selection of flies, based on minimum number of landmark visits, are noted in the figure. Below each boxplot, the total number of visits to any landmark (median across flies) is shown as a heat map (color code in F, right side).

### Pairing landmark shapes with virtual sugar increases non-specific landmark attraction

We asked whether a fly’s naïve landmark shape preference could also be modified by selectively associating one landmark shape with *virtual sugar* stimulation. Because *virtual sugar* may partly —but incompletely— mimic an appetitive reward, we expected that flies might respond to such an experience by selectively increasing their interaction with the ‘rewarded’ landmarks. As observed previously in the local search experiments, flies slowed down upon exposure to the virtual sugar (**Fig. S7G** center: increased residency around cones). Their altered walking behavior also led to increased visits and interaction with landmarks paired with *virtual sugar* stimulation —as expected from the local search experiments— but this effect did not persist in form of a selective increase in approaches or visits to the ‘rewarded’ landmark once the *virtual sugar* stimulation ceased (**Fig. S7H,I** compare Training and Post trials). Rather, flies showed an increase in landmark visits to both landmarks (**Fig. S7H,I** total visit count in Pre and Post trials). Thus, *virtual sugar* reward is either insufficient to mimic key features of food reward consumption, or appetitive conditioning with landmarks requires color or other, stronger differences between their appearance.

## Discussion

We combined 2D visual VR and optogenetic stimulation to study how head-fixed flies change their behavior upon experiencing sensations of sugar and heat. Head-fixed flies respond to transient encounters with *virtual sugar* by changing their walking patterns much as free walking flies do —by exhibiting a local search behavior. In free-walking flies, this search behavior has been suggested to rely on path integration [27], but this was not clear for flies in our tethered setting. The most prominent behavioral change we observed could also be explained by simpler mechanisms, such as a modulation of the rate and tightness of turns [58-60]. We were unable to use the appetitive experience of virtual sugar to differentially condition flies to specific visual landmarks. However, we found that location-specific *virtual heat* was sufficient to condition head-fixed flies to alter their landmark preferences.

Although visual place learning has been demonstrated in freely walking flies in 2D environments [43], studies of visual conditioning in head-fixed flies have relied on 1D environments in which flies are trained to avoid aversive heat by orienting toward certain visual cues [61]. Our paradigm relied on optogenetically stimulating heat sensing neurons, but the detailed knowledge of —and genetic access to— sensory and reinforcement pathways in *Drosophila* should permit the generation of other “virtual” sensory stimuli using optogenetics [35, 62, 63]. Using optogenetically-generated virtual stimuli offers significant advantages for experimental design by enabling the creation of sensations in flies that might otherwise affect their state of health (for example, heat, which can physically damage a fly’s body, or sugar, which might satiate it). This experimental approach also allows different sensory modalities to be flexibly paired with the visual VR without the need of hardware or software modifications. Such flexibility could permit screens in which specific pathways can be activated in isolation to identify the role of different cell types. In addition, the time course of a *virtual heat* or *sugar* stimulus is easier to control, and does not rule out the use of thermogenetics to modify the activity of different neural populations [64-66].

Our operant visual conditioning paradigm required a head-fixed animal to sample its virtual environment, learn how the unconditioned stimulus related to the visual environment, and alter its behavior based on this relationship within 20 minutes of training. Operant learning is expected to be harder in 2D than in angular 1D environments, because, unlike in 1D, the visual scene associated with any given reinforced location is not unique and depends on the fly’s heading. Fully sampling this large space of visual stimuli paired with reinforcement requires time, making the task of learning the association significantly more challenging. This is true even in insects that are well known to rely on visual learning in natural settings [67]. Training flies for longer time periods or biasing the sampling of the environment during training may further strengthen conditioning. Moreover, optogenetically generated virtual heat is likely to be experienced differently than real heat: Flies possess multiple heat sensors feeding into distinct temperature processing pathways [68], many of which may be activated in real heat reinforcement used in existing visual conditioning assays. Nevertheless, our results suggest that activation of HC neurons and their downstream partners is sufficient to induce a visual memory.

What exactly the fly learns during operant visual conditioning is an open question. Based on visual conditioning experiments in 1D visual environments [61] and observations in other free-behaving insects [69, 70], it has been proposed that insects use a snapshot-based visual learning mechanism. The idea of snapshot learning is that the animal associates a specific image template on its retina with the reinforcement rather than a more generally recognizable feature. Later studies suggested that flies can generalize learned associations with patterns to some degree [71], which would be advantageous for visual learning in 2D environments where the same visual landmark can generate a variety of different visual stimuli on the retina depending on the fly’s heading direction. In 2D environments there may also be multiple valid behavioral adaptations within a given conditioning paradigm. In our paradigm, for example, flies could learn to selectively approach the non-punished landmark or avoid the punished one. Furthermore, these adaptations may be specific to certain locations within the environment: as an animal approaches a landmark, naïve preferences for landmark fixation might be able to override a learned aversion as the image of the looming landmark becomes more salient. 2D VR provides a tool for probing these interactions in more detail.

In mammals, tracking animals moving in 2D or 3D settings while simultaneously recording from their brains has uncovered navigational strategies and neural representations thought to be involved in goal-directed navigation [72]. In smaller animals, however, neural recordings at cellular resolution require the animal to be head-fixed, which poses a challenge for studying the neural basis of navigational behaviors. Our 2D VR paradigms should make it possible to study the behavior of head-fixed flies under conditions that accurately capture features of visual stimuli typically present during open-field navigation, such as optic flow and looming during translational motion. We specifically designed our VR system and behavioral paradigms to be compatible with two-photon calcium imaging, which should enable future investigations of neural dynamics underlying goal-directed navigation in 2D environments.

## Acknowledgments

We thank Brian Coop and Anthony Leonardo for help with building the projector-based display; Albert Lee for sharing the MouseoVeR code base as a basis for the FlyoVeR software; Laura Porta (L.P.) for pilot experiments with Gr64f; Abel Corver for help with the Gr64f conditioning experiments; TJ Florence, Ed Rogers and Gabriella Sterne for sharing flies from their local stocks; and Michael Reiser, Eugenia Chiappe, Rebecca Yang, TJ Florence, Yoshi Aso, Tilman Triphan, Claire Eschbach, Alice Robie, Ann Hermundstad and members of the Jayaraman lab for discussions and advice. We thank Janelia’s Project Technical Resources team for dissections, immunolabelling and imaging to confirm expression patterns. We are grateful to members of the Jayaraman lab for feedback on the manuscript. B.A., M.A.B. and L.P. were supported by the Janelia Undergraduate Scholar program. This work was supported by the Howard Hughes Medical Institute.

## Contributions

H.H. and V.J. designed the project. B.A. built a pilot VR setup with help from J.D.C. M.B., and C.B. developed the FlyoVeR software with input from H.H., J.D.C., and V.J. H.H., with input from V.J., built the VR setup and wrote software for hardware control. H.H. designed virtual worlds, built the free walking arena, and performed optogenetically-triggered local search, virtual heat avoidance and conditioning experiments. H.H. and M.A.B. performed crosses, prepared flies and performed fixation behavior experiments. H.H. analyzed all data. H.H. and V.J. wrote the manuscript

---

**Supplementary Figure 1:**
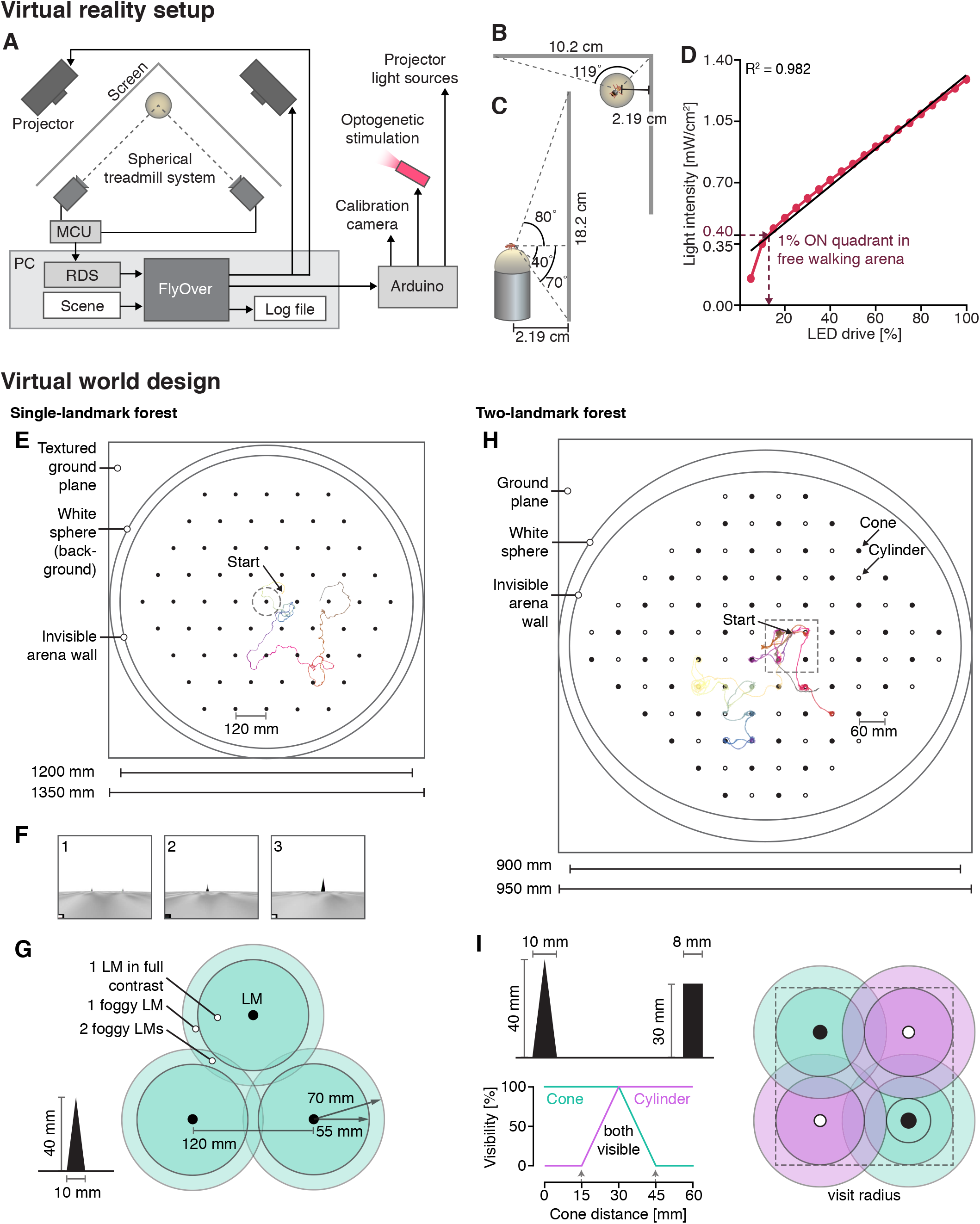
Design of the 2D VR system. (**A-D**) Extended information on the hardware and software of the VR system. (**A**) Schematic of the hardware and software control loops. (**B, C**) Schematic illustrating the size of the visual display screen and its position relative to the fly on the treadmill. View from the side (C) and top (B). The range of the horizontal field of view indicated in (C) corresponds to the closest screen distance relative to the fly, which occurs at 90° on both sides. The virtual scene horizon is positioned at the height of the ball (dashed horizontal line) (**D**) Light intensity measured on the ball surface with a power meter (Methods) for different input LED drives (corresponding directly to different reinforcement levels). Except for low light intensities, the relationship between LED drive and light intensity was well described by a linear fit (y = 0.05 × + 0.26, R^2^ = 0.98). (**E-I**) Design of the periodic virtual worlds. (**E**) Schematic of the complete *single-landmark forest* scene with an example trajectory from a 10 min trial overlaid. (**F**) Image frames from different time points during the approach of a landmark. Note that, in contrast to the frames in Fig. 1, these are screen shots taken from the actual panorama that was projected onto the screen, thus reflecting the distortion that was used to account for the screen geometry. (**G**) In the periodic world design, landmarks are positioned on the nodes of equilateral triangles that form the unit cell of a large hexagonal grid. The shortest distance between two adjacent landmarks is 120 mm. The two shaded circles around each landmark indicate the where the respective landmark begins to be visible (lightly shaded circle: 70 mm radial distance) and where it starts to appear in full contrast (darker shaded circle: 55 mm radial distance). Cone-shaped landmarks were 10 mm wide at the base and 40 mm tall. (**H**) Schematic of the *two-landmark forest* scene with an example trajectory from a male HC-Gal4 > ChrimsonR fly (10 min pre-trial, group trained against the cylinder; Methods). The dashed circle in (E) and the square in (H) indicate the “unit cells” onto which walking trajectories through the periodic scene are projected. Note the different spatial scales in the schematics in (E) and (H). (**I**) Schematic illustrating landmark placement and dimensions as well as virtual fog settings in the *two-landmark forest* scene. Graphic on the right illustrates visibility with shaded circles around each landmark analogously to (G). RDS, remote data server; MCU, microcontroller unit; LM, landmark.

**Supplementary Figure 2:**
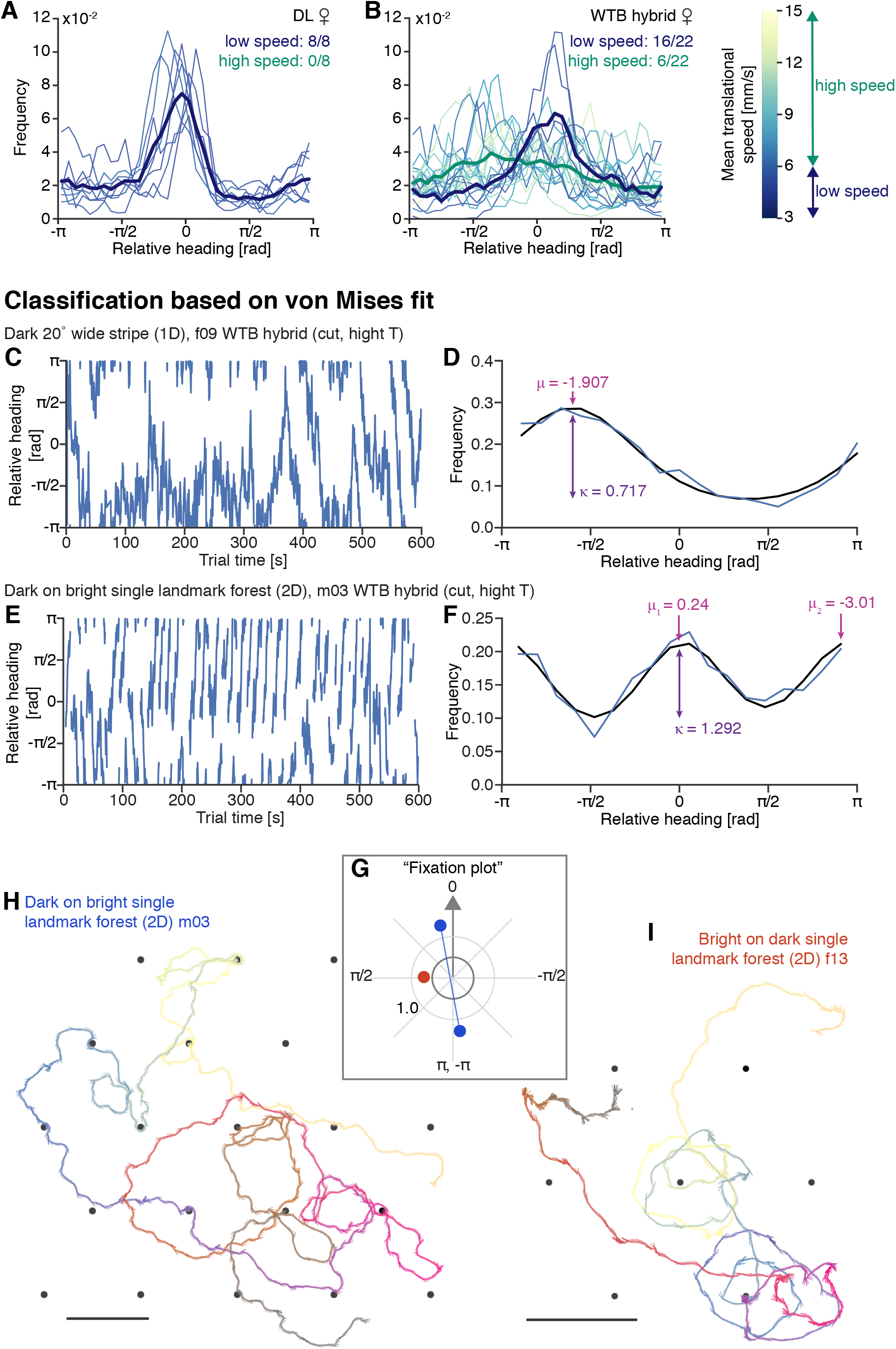
Extended data on stripe and landmark fixation. (**A, B**) Effect of walking speed on fixation as quantified in this paper. Relative heading distributions calculated from data of female DL flies (A) and female WTB flies (B) after separating flies into two groups based on their mean translations speed computed over the full trial (criteria illustrated in (B), left). The number of flies that fell into each group is indicated in the figure. (**C-I**) Classification of fixation behavior based on von Mises fit. (**C,E**) Time series of the relative heading angle of two female WTB flies (cut wings, 30 °C room temperature) over a 10 min trial with either a dark 20 ° wide stripe on bright background (C) or in a dark 2D plane with bright landmarks (E). (**D,F**) Blue line: Relative heading distributions computed from the time series shown in (C, E), respectively. Black line: von Mises fit to the measured distribution. In (D) a unimodal and in (F) a bimodal von Mises distribution was fitted. The location (μ) and shape (κ) parameters are indicated in the plot. See Methods for details on the fitting procedure. (**G**) Polar "fixation plot” visualizing fitted von Mises parameter for the two fixation trials shown in (H, blue) and (I, red). (**H,I**) Walking traces of flies in a 2D landmark fixation trial. Progression of time is color-coded as in Fig. 1H. Scale bar: 100 mm. Panel (H) shows data from same fly as (E,F).

**Supplementary Figure 3:**
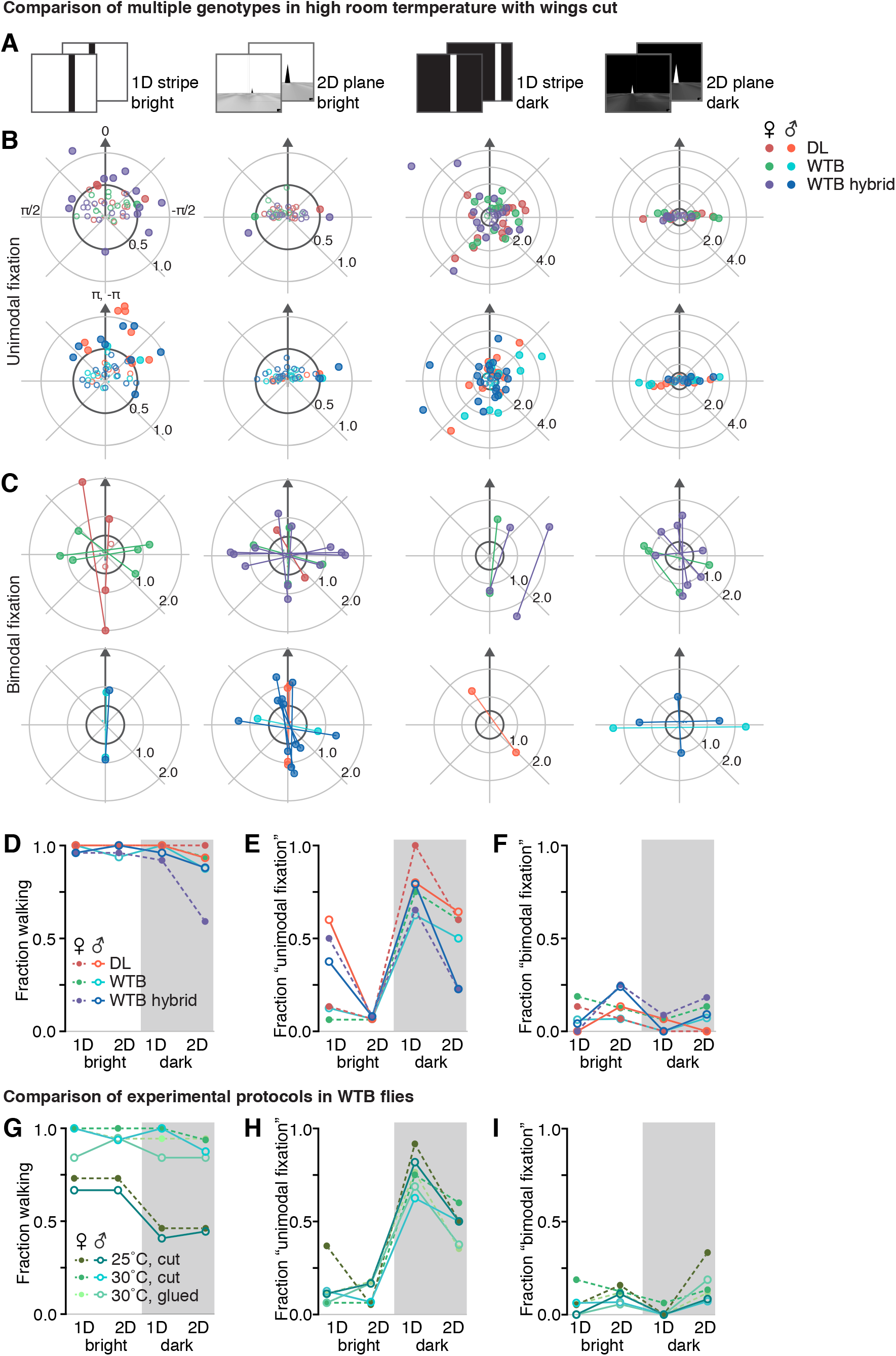
Fixation behavior across sexes and genotypes. (A-C) Comparison of fixation behavior in WTB, WTB hybrid and DL flies measured at high room temperature (30 °C) and with wings cut. (**A**) Illustration of the four trials used to test the effect of different contrast conditions. (**B**) Fixation plots of unimodal fixation fits of female (top) and male (bottom) flies. Visualization as explained in Fig 2. (**C**) Fixation plots of bimodal fixation fits of female (top) and male (bottom) flies. Note different axis scale in (B,C). (**D**) Fraction of flies that walked for at least 20 % of the trial time across the four trials. Data from male and female flies of three genotypes color-coded as indicated in (B, top right). (**E,F**) Fraction of flies showing unimodal (E) and bimodal (F) fixation. (**G-I**) Same as (D-F), but for different experimental protocols: Flies were either wing cut and measured at high (30 °C) or low (25 °C) room temperature or wing glued and measured at high (30 °C) room temperature.

**Supplementary Figure 4:**
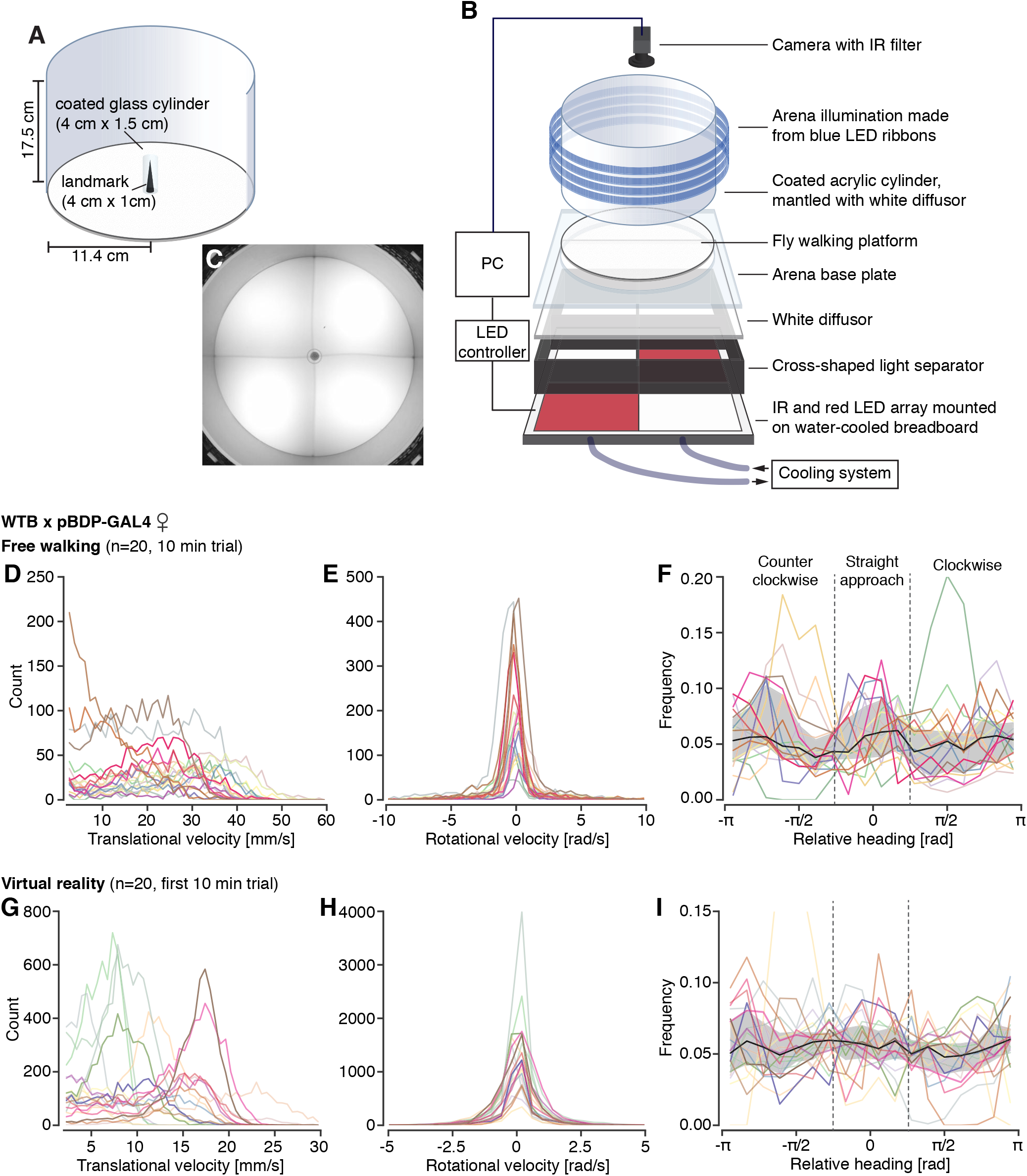
Landmark interaction in freely and tethered walking flies. (**A**) Schematic of the free walking arena with a landmark and the glass cylinder that prevents flies from climbing onto the landmark. (**B**) Schematic of the entire free walking rig. (**C**) Video frame from a trial with a male fly (visible in upper right arena quadrant) exploring the arena with a landmark. The image was taken with an IR camera and does not reflect the lighting conditions visible to the fly. (**D-I**) Data from female WTB hybrid flies free walking trials (**D-F**, n = 20) and VR trials (**G-I**, n = 20, only first out of three VR trials with visible landmarks included). Fly identity is color-coded. (**D**,**G**) Histograms of the translational velocity. (**E**,**H**) Histogram of the rotational velocity. (**F**, **I**) Relative heading distributions, visualized as in Fig. 2A. Relative heading angle bin size: 20 °. For free walking data, only trajectories within 10 mm – 60 mm radial distance from the landmark (center of the arena) were considered in the analysis.

**Supplementary Figure 5:**
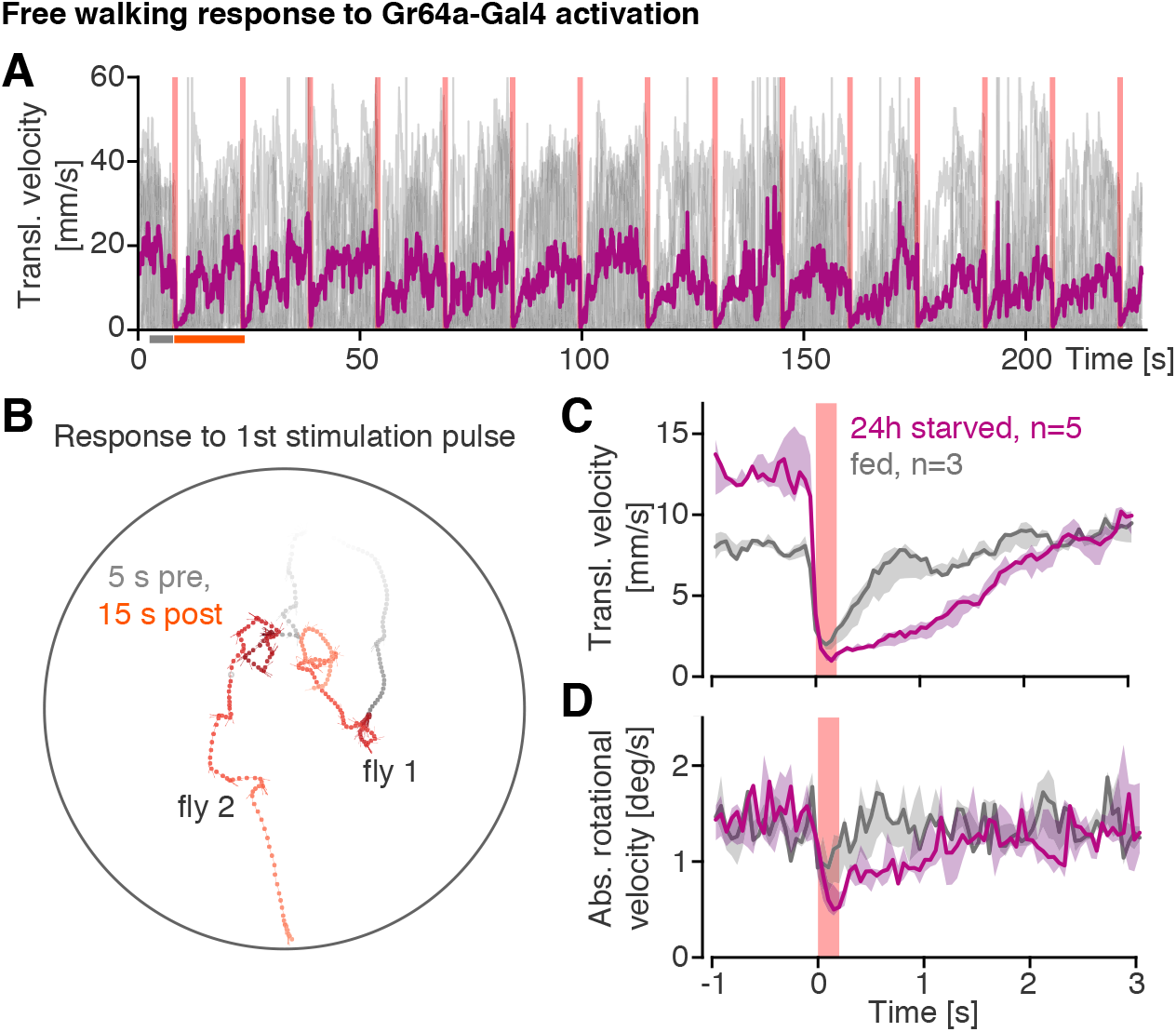
Effects of optogenetic activation of sugar sensing neurons. (**A**-**D**) Stopping response to optogenetic activation of sugar sensing neurons in freely walking flies. (**A**) Translational walking velocity of female Gr64f > ChrimsonR flies (24 h starved) over the time course of the full protocol consisting of 15 light stimulation pulses (200 ms, 1.58 mW/cm2) each separated by a 15 s break. Walking behavior was measured in groups of ~10 flies. Grey lines: velocity traces from individual flies. Purple line: Mean translational velocity across flies. (**B**) Two example trajectories from the first stimulation pulse. Trajectories are colored in grey before the stimulation (5 s) and in red after (15 s). Circle marks arena border. (**C,D**) Average response of 24 h starved (purple) and fed (grey) flies to a stimulation pulse. The responses in translational (C) and rotational (D) walking velocity are shown as averages (median and interquartile range) across experimental repeats (starved group: n = 5, fed group: n=3). For each repeat, responses were averaged (mean) across flies and subsequently across the 15 stimulus repetitions.

**Supplementary Figure 6:**
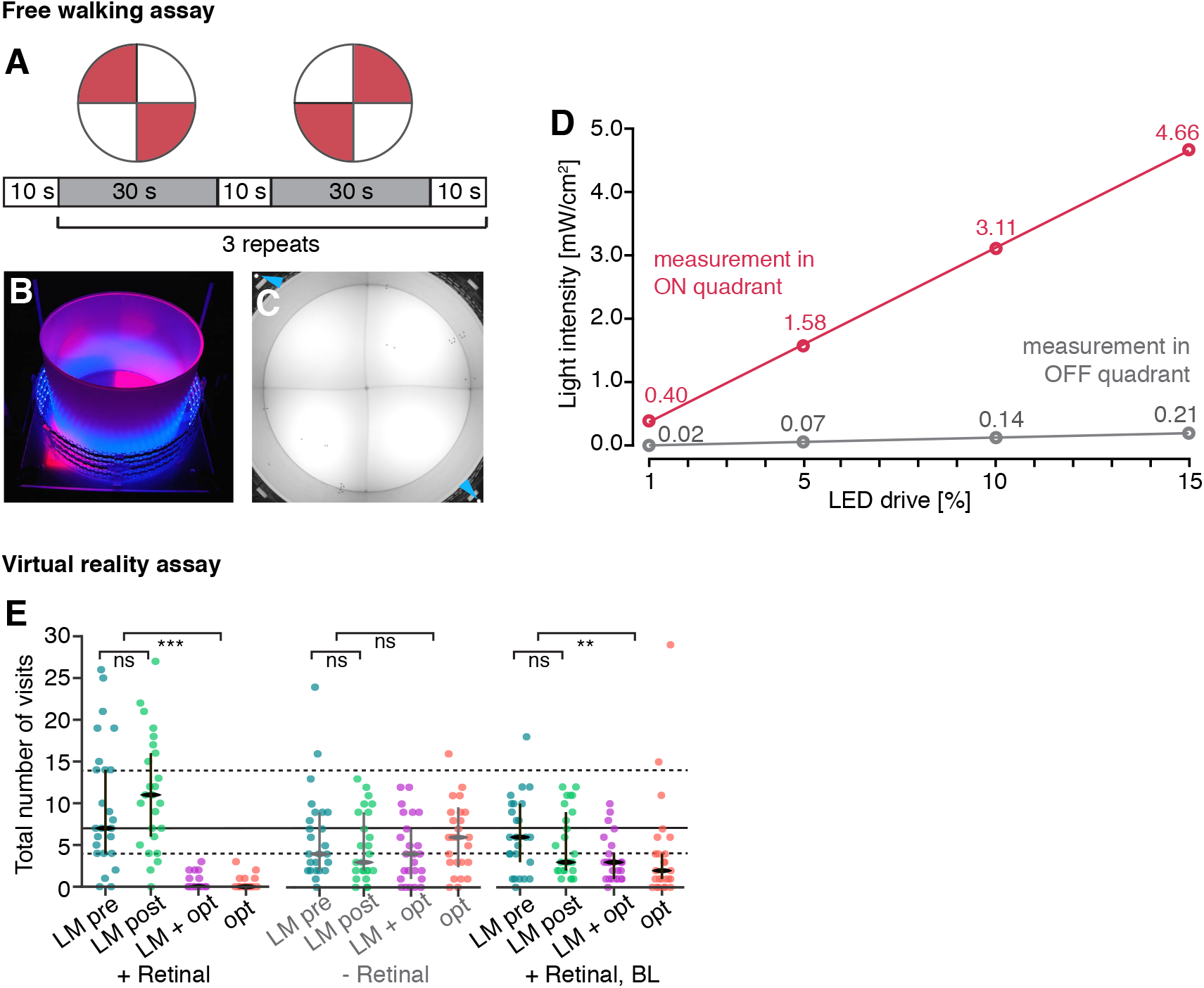
“*Virtual heat*” avoidance in free walking and VR. **(A-D)** “*Virtual heat*” avoidance in **(A)** Schematic of the quadrant assay trial structure. **(B)** Photograph showing the lighting conditions during the quadrant assay. Blue LEDs around the arena wall provide ambient light visible to flies. The red light illuminating two quadrants is invisible to flies. **(C)** One frame from a video recorded during a trial with male HC-GAL4 × 10xUAS-ChrimsonR flies. Two IR lights at the corners of the image (arrowheads) indicate which quadrants are illuminated. In this frame all flies have moved out of the illuminated quadrants. **(D)** Measured light intensities as a function of % LED driver current in the illuminated “ON” (red points) and the not illuminated “OFF” quadrants (grey points). Measurements were performed during continuous light illumination with a power meter (see Methods). The solid lines are the linear regressions for the five measurements. ON quadrants: y = 0.30x + 0.08, R^2^ = OFF quadrants: y = 0.01x - 0.00, R^2^ = 1.00. **(E)** *Virtual heat* avoidance in VR. Comparison of the total number of visits across trials and three experimental groups. Dots mark measurements from single flies, with Median and IQR are overlaid in black or grey, respectively. The solid and dashed line serve as a reference for the visit count in the first landmark only trial in retinal-fed flies with no virtual heat baseline. We used a paired Wilcoxon test to compare the total visit count between the pooled trials with and without optogenetic stimulation, and the “LM pre” and the “LM post” trials of each group. Significance codes: “ns” p > 0.1, “*” p ≤ 0.05, “**” p ≤ 0.01, “***” p ≤ 0.001. LM, landmark; opt, optogenetic stimulation; r_z_, reinforcene radius, BL, baseline.

**Supplementary Figure 7:**
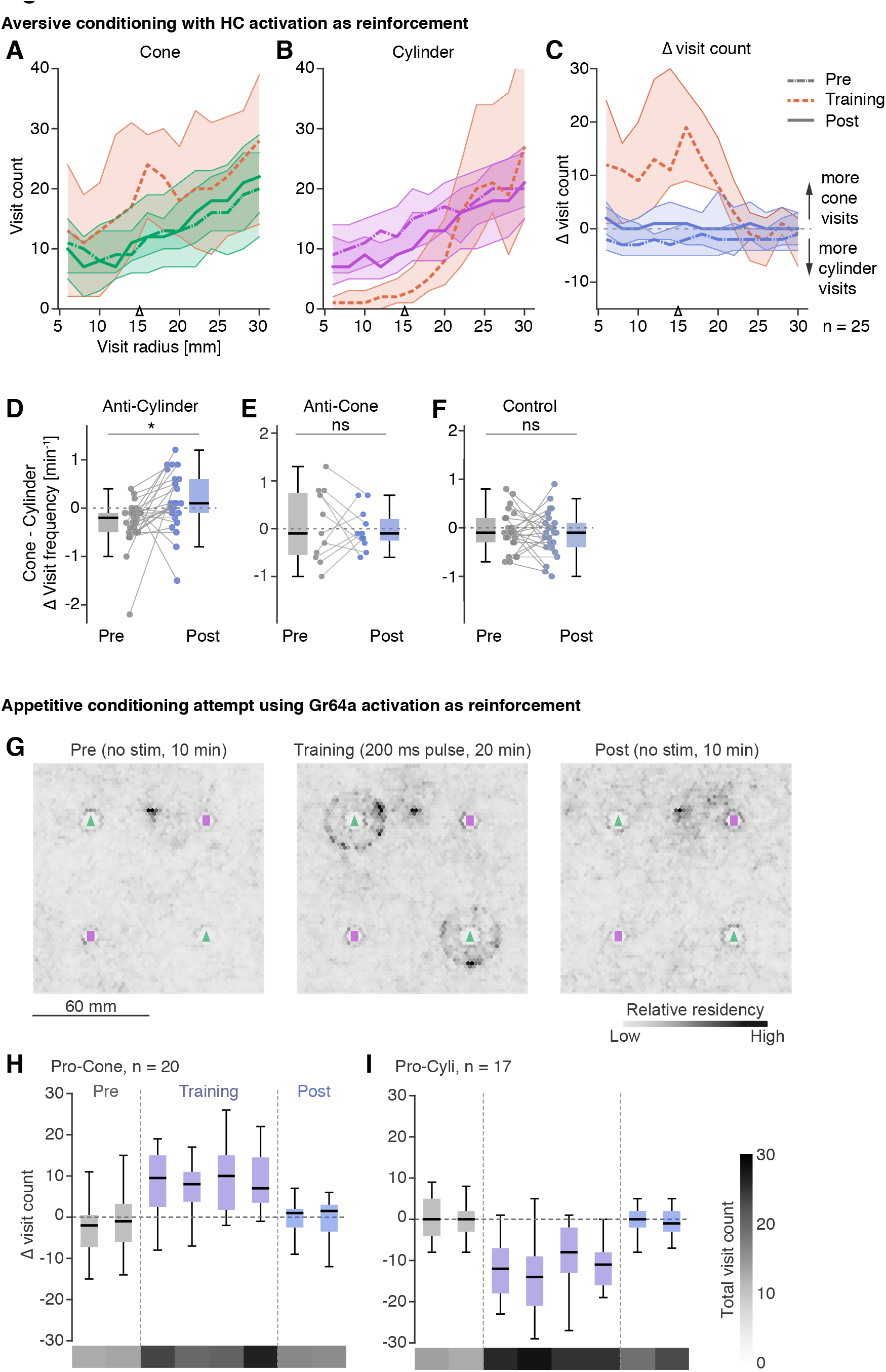
Extended data on conditioning with “virtual heat”. (**A, B**) Visit counts for cone (A) and cylinder (B) as a function of visit radius. Median (thick line) and IQR (shaded region) are shown for the three trials. Line style of the median encodes trial identity (see legend in C, right). (**C**) Difference in visits (cone visits – cylinder visits) as a function of visit radius. Visualization as in (A,B). (**D**-**F**) Delta (∆) visit frequency computed as cone visit frequency - cylinder visit frequency in the pre- and post-trial for the three conditioning protocols presented in Fig. 7 (E, F, G). Positive values correspond to a relative preference for visiting cones. For each group, we performed a paired t-test with for alternative hypothesis that the true location shift is 0. Anti-cylinder training (D): n=22, p=0.00305 (*), anti-cone training (E): n=11, p=0.74895 (ns), control training (F): n=22, p=0.56107 (ns). (**G-I**) Attempted appetitive conditioning based on Gr64f-Gal4 neuron activation in the *two-landmark forest* VR environment. (**G**) Residency distribution visualized as in Fig 2E for the three trials of the conditioning paradigm: a pre trial (10 min) with no optogenetic stimulation, a 20 min training trial with stimulation around cones (20 mm radius) and a 10 min post trial, again with no stimulation. (**H,I**) Comparison of shift in landmark preference across two experimental groups: pro-cone training (H, same as in G) and pro-cylinder (I) Landmark preference was quantified for each fly in intervals of 5 min as the number of cone visits minus the number of cylinder visits. Data from all flies that made at least 5 visits to any landmark in each trial is presented as boxplots (black line: median, box spans the 25th to the 75th quartile). Sample sizes after selection of flies, based on minimum number of landmark visits, are noted in the figure. Below each boxplot, the total number of visits to any landmark (median across flies) is shown as a heat map (color code in I, right side). Significance codes: “ns” p > 0.1, “*” p ≤ 0.05, “**” p ≤ 0.01, “***” p ≤ 0.001.

## Supplementary Movies

<u>Supplementary Movie 1:</u> ***Landmark preference experiment***

(**top left**) Trajectory of a male WTB hybrid fly walking through the *two-landmark forest* VR. Landmark locations are marked by grey triangles (cone positions) and rectangles (cylinder positions). The colored dot indicates the fly’s virtual position, with the black stick pointing in the fly’s heading direction. The color of the dot changes with progression of time. Past positions of the fly in VR are marked by small dots. (**bottom left**) Collapsed version of the trajectory. (**top right**) Visual scene as seen from the fly’s point of view. Note that images were rendered with 120 degree FOV rather than 240 degrees used in experiments. Also, images shown here were not perspective corrected as would be for the fly. (**bottom right**) Video of the tethered fly walking on the spherical treadmill. The image on the panoramic screen can be seen in the background.

<u>Supplementary Movie 2:</u> ***Aversive visual conditioning experiment***

Illustration of the walking trajectory of a male HC-Gal4 > ChrimsonR fly during training in the aversive visual conditioning paradigm. (**top**) The large colored dot indicates the fly’s position in the virtual world. The fly’s position is visualized as in Movie 1, except that here the color shows the optogenetically induced *virtual heat* level at the respective position, and object identity is also color-coded (same color code as in Fig. 5b). (**bottom**) Visual scene as seen from the fly’s point of view. As in Movie 1, images were rendered with 120° FOV rather than 240° used in experiments and not perspective corrected as would be for the fly.

## Methods

### EXPERIMENTAL MODEL AND SUBJECT DETAILS

#### Fly strains

Three wild type strains were used: the isogenic wild type Berlin (WTB, donated by the Heberlein lab, [73]) strain, the Dickinson laboratory (DL, gift from the Reiser lab, [74]) strain and WTB hybrid flies. WTB hybrid flies were generated by crossing WTB virgins with males from an enhancerless “empty GAL4” line (pBDPGAL4U in attP2, [75]). We chose WTB flies because this genotype has been used in many previous publications on various walking behaviors [76, 77]. WTB hybrid flies were chosen to approximate the genotypes used in optogenetic activation experiments, where an effector line with WTB background is crossed to GAL4 driver lines with variable genetic backgrounds. For optogenetic stimulation experiments we crossed either Gr64f-Gal4 [53] or Hot cell (HC)-GAL4 [57] to 10xUAS-ChrimsonR-mVenus flies (trafficked in VK00005, WTB background, generated at Janelia Research Campus). The Gr64f-Gal4 line (w; Gr64f-GAL4(737-5)/CyO; Gr64f-GAL4(737-1)/TM3) was constructed from <u>w^*^; P{Gr64f-GAL4.9.7}5/CyO; MKRS/TM2</u> (Bloomington Stock Center, stock #57669) and w^*^; <u>P{UAS-mCD8::GFP.L}LL5</u>; <u>P{Gr64f-GAL4.9.7}1/TM3</u>, Sb^1^ (Bloomington Stock Center, stock #<u>57668</u>).To confirm expression patterns of HC-Gal4 we used two driver lines: pJFRC2-10XUAS-IVS-mCD8::GFP and pJFRC12-10XUAS-IVS-myr::GFP (both Janelia Research Campus).

#### Fly rearing conditions

Flies were reared at 23 °C in 60 % relative humidity with a 16:8 light:dark cycle on fly food that was prepared according to a recipe from the University of Würzburg, Germany. Crosses for generating flies expressing ChrimsonR [78] were set on Würzburg food with added all-*trans*-retinal (0.2 mM concentration) and offspring were transferred onto food with a higher retinal concentration (0.4 mM). Flies expressing the opsin were reared in low-light condition in a blue acrylic case inside the incubator to prevent: (a) activation of ChrimsonR-expressing neurons and (b) degradation of retinal in the food. We kept flies under low-light rather than in darkness to expose the flies to visual stimuli prior to behavioral experiments, and to ensure a normal circadian rhythm. Control flies for optogenetic activation experiments were also reared inside blue cases, but on standard Würzburg food. Cornmeal and molasses in the standard Würzburg food are a potential source for all-*trans*-retinal [79, 80]. The exact retinal content of the standard food may vary, but we expect it to be well below 0.001 mM.

### METHOD DETAILS

#### Confirmation of expression patterns

We used HC-Gal4 > mCD8::GFP (4-6 d) and HC-Gal4 > myr::GFP (6-7 d) to confirm the expression pattern of HC-GAL4. Dissections, immunolabelling and imaging were performed as described in [81]. We looked at the expression pattern in the brain (mCD8::GFP: 7 female, 5 male; myr::GFP: 4 female, 5 male) and the ventral nerve chord (VNC, mCD8::GFP: 5 female, 4 male; myr::GFP: 4 female, 4 male). The image shown is a montage of maximum intensity projections of two-color stacks of the brain and the VNC.

#### Preparation of flies for behavior experiments

Prior to experiments, 3-5 day old flies were cold anesthetized, sorted by sex and the distal two thirds of their wings clipped. The decision to use wing-clipped flies was motivated by two observations. Firstly, clipping the wings 1-2 days prior to experiments strongly reduced the rate of attempted take-offs or jumps of tethered flies on the ball. Secondly, many previous studies on visual navigation in flies have used wing-clipped flies (for example, [76]), making it possible to compare our data to those results. After wing-clipping, male and female flies were kept separately and transferred into fresh food vials with a small piece of filter paper, where they were allowed to recover for 2-3 days before experiments. Experiments were performed with 5-10 day old flies. For some experiments, we used an alternative technique to render flies flightless: gluing the wings together in a relaxed position with a small drop of glue right behind the thorax. For experiments in VR, wing-clipped flies were cold-anesthetized and glued to a thin tungsten wire pin with UV-curable glue (KOA 300, KEMXERT, York, PA, USA). With an additional small droplet of glue, deposited above the neck connective, the head was fixed to the thorax, keeping it in a relaxed position. We decided to fix the head to minimize movements of the fly that would disrupt closed-loop visual stimulation. For experiments with *virtual sugar* stimulation, flies were wet-starved for 24 h prior to experiments by transferring wing-clipped flies to vials containing only a humidified piece of filter paper. Such wet starved flies were then tethered to a pin and used in behavioral experiments within 2 h (effective starvation 24-26 h). For all other experiments, flies were taken directly out of the food vials and tethered to the pin, but were then transferred to VR rig and tested within 3-6 h. Flies were given 10-30 min to adjust to the ball before starting an experiment.

#### 2D visual VR

##### Miscellaneous hardware

On the ball, the fly was surrounded by a triangular screen formed by two 18.2 cm high and 10.2 cm wide display screens (**Fig. 1A** and **Fig. S1A-C**). The fly was located symmetrically between the two screens, 3.1 cm behind the tip of the triangle and at one third of the screen height. The distance between the fly and the screens varied along the azimuthal direction and was smallest at 90° to either side with 2.19 cm. The composite screen spanned 119° of the fly’s azimuthal field of view (FOV) on both sides. The coverage along the vertical FOV was limited by the ball at the lower edge to 40° below the horizon line. The vertical coverage above the horizon line varied with the distance of the fly from the screen being highest where the fly was closest to the screen with 80° and lowest at the two tips of the screen with 57°. The screen was made from a single white diffuser sheet (V-HHDE-PM06-S01-D01 sample, BrightView Technologies, Durham, NC, USA) that was enforced around the edges, folded along the middle and mounted onto a custom-made metal frame. The projectors were connected to a computer via two display ports on a graphics card (GeForce GTX 770, Nvidia, Santa Clara, CA, USA) with three independent outputs. For closed-loop optogenetic stimulation in VR, we used a red LED directed at the fly on the ball (625 nm, M625F1, Thorlabs Inc, Newton, NJ, USA). We calibrated the optogenetic stimulus by measuring the light intensity with a power meter (PM100D with S130C Sensor, at 625 nm with range setting 1.3 mW) for a range of LED drives (**Fig. S1D**). We adjusted the room temperature and humidity to maintain a temperature of around 28-30 °C and a relative humidity of 28-32 % in the VR rig.

##### Spherical treadmill

We used a spherical treadmill as described in [22]. The treadmill ball was hand-milled from polyurethane foam (FR-7120, Last-A-Foam, General Plastics Manufacturing Company, Tacoma, WA, USA) and had a diameter of 9.93 mm and a weight of 37 mg. The ball was freely floating on an aircushion in a custom-made holder. The airflow to the ball was maintained at 0.45 l/min using a mass flow controller (Alicat Scientific, Tucson, AZ, USA) and humidified by passing it through a bottle humidifier (Salter Labs, Lake Forest, IL, USA). The ball surface was illuminated from below and the side with a set of four IR LEDs emphasizing the texture of the ball surface for high tracking performance. The ball motion was captured by a previously described ball tracker system [22]. The treadmill readout had arbitrary spatial units per time (au/s). To obtain a ball rotation velocity measurement in mm/s the treadmill output was calibrated using a third (calibration) camera and the same calibration procedure as described in [22]. We used a custom MATLAB script and a programmable microcontroller (Arduino Mega 2000, **Fig. S1A**) to generate a trigger sequence for the calibration camera that was synchronized with a treadmill recording.

##### Projector-based visual display

The visual display consisted of a triangular screen onto which a panoramic image was back-projected by two DLP (Digital Light Processing, also referred to as digital mirror device or DMD) projectors (DepthQ WXGA 360 HD 3D Projector, developed by Anthony Leonardo, Janelia, and Lightspeed Design, Bellevue, WA, USA). The two projectors each generated an image with 720 × 1280 pixel resolution, and were aligned to generate a continuous panorama (1440 × 1280 pixel). At the closest point (90° to either side, 2.19 cm distance), a pixel subtended a visual angle of 0.74°. The furthest pixels on the screen, located in the upper back corners of the screen (10.65 cm distance), subtended an angle of 0.15°. Thus, the maximum angular pixel size in our setup is well below 5°, the approximate interommatidial angle of *Drosophila melanogaster* [82], ensuring that the movement of images across the screen appears smooth to the fly. The projectors we used were customized to deliver visual stimuli to the fly. The color-wheel was removed to allow the display of 8-bit grey-scale images at a frame rate of 360 Hz. This ensured a refresh-rate above the animal’s flicker fusion frequency, making the projected movements look continuous to a fly [83-85]. Furthermore, the optics were optimized for close-range projection and the lamp was replaced with a light guide to a blue LED light source (458 nm wavelength, SugarCUBE LED Illuminator, Edmund Optics Inc, Barrington, NJ, USA). Although blue light is not the ideal choice for behavioral assays, we chose this wavelength to permit easier transfer of the behavioral paradigm to commonly used two-photon calcium imaging setups in the future. We measured the irradiance of the projected image on the panoramic screen with a power meter (PM100D with S130C Sensor, Thorlabs Inc, Newton, NJ, USA, sensor facing toward the projector) for a wavelength of 459 nm (peak wavelength in previously measured spectrogram). When a bright scene with one dark landmark was projected onto the screen, we measured a light intensity of 0.52 mW/cm^2^ in the center of the right screen and 0.54 mW/cm^2^ on the left screen. The light intensity within the image of a black landmark projected onto the center of the right screen was 0.02 mW/cm^2^. Thus, the projected image had a Michelson contrast of about 0.93.

##### Software for 2D VR

The treadmill system tracked the ball’s movements at 4 kHz and this data was downsampled to 400 Hz by a custom-written C++ application (Remote Data Server, RDS) and passed on to FlyoVeR. The FlyoVeR application is a modified version of the Jovian/MouseoVeR software [45] (see www.flyfizz.org for details). We achieved a high refresh rate of the displayed images by rendering sets of three frames at a time in FlyoVeR and packing them into the red, green and blue color channel for a full color frame buffer, which were displayed sequentially and refreshed at 120 Hz. FlyoVeR computed the fly’s walking velocities from the treadmill’s ball rotation measurements as described in [22]. The positions were then integrated to compute the fly’s position. The two calibration factors that are necessary to convert the treadmill output from [au/s] to [mm/s] (see www.flyfizz.org for details) as well as the ball radius were provided through a GUI. A small square at the edge of the projected image, whose color was toggled between white and black with each newly drawn frame, served as a frame rate indicator, which could be read out with a photodiode. Data on the fly’s virtual position and velocity was logged at 360 Hz.

FlyoVeR also sent a reduced output stream (at 60 Hz) via a serial connection to a microcontroller (Arduino Mega 2000). The microcontroller then set the red stimulation LED to the light intensity specified by the reinforcement level parameter. We measured the relationship between current input (characterized in % of maximum current) and light intensity for the red stimulation light, which was, with the exception of very low input currents, approximately linear (**Fig. S1D**). We used this empirical calibration curve to translate light intensities that worked in the free walking arena to light levels to be used in VR.

#### Virtual world design

Custom 3D scenes were designed in created with the free 3D modeling program Blender (version 2.73) and were exported in Collada (version 2.4) format, a standardized XML format used to describe 3D graphics. Collada files could then be loaded into FlyoVeR though the graphical user interface.

Each object within the 3D scene had a unique name and a set of properties. Properties such as the color and texture were specified in Blender as “materials” that were then assigned to the respective object. Other properties such as object visibility were communicated to FlyoVeR as part of the object’s name string (see www.flyfizz.org for details).

##### 1D stripe fixation

For 1D stripe fixation experiments, we generated a 3D scene consisting of a cylinder centered around the fly’s virtual position. The cylinder was subdivided into 360 vertical faces, such that each face corresponded to a vertical strip of 1° angular width when seen from the cylinder center. For dark-on-bright stripe fixation experiments we used a cylinder with 20 consecutive faces colored black and all other faces white, corresponding to a 20° wide black stripe on a white background. For the *bright-on-dark* stripe fixation experiments the colors were reversed.

##### 2D scenes

Virtual worlds for 2D VR experiments consisted of a large textured ground plane with sparsely distributed landmarks (**Fig. S1E,H**). In most experiments, we used a scene with black landmarks on a bright background and a lightly textured ground plane, matching the conditions in the free walking assay (dark-on-bright condition). For the ground plane texture, we chose a white-noise grey scale pattern with pixels varying either between 0 % and 50 % black level (low contrast, see images in **Fig. S1F**) or between 0 % and 100 % (high contrast). The low-contrast ground plane was used in all experiments, except when making comparisons to free walking. Landmarks were always impenetrable, that is, the fly could not move into or through the virtual landmarks. The dimensions of the virtual landmarks were chosen to match real landmarks that induced frequent approaches in freely walking flies (data from a pilot study). Free walking experiments also informed the spacing of the virtual landmarks relative to each other. The area of the virtual world that flies could explore was bounded by an invisible, impenetrable cylinder (**Fig. S1E,H**). A white, flattened sphere encompassing the arena border and the ground plane served as a backdrop. The overall size of the virtual world was chosen based on pilot experiments in VR to ensure that most flies did not reach the arena border within a 10 min trial. Each visible landmark was surrounded by a slightly (0.5 mm) larger invisible object of the same shape to prevent visual artifacts when a fly came close to the surface of the landmark. In bright-on-dark trials the color of all components was inverted, i.e., we used white landmarks, black fog and the ground plane texture pattern was inverted.

We used two types of 2D scene geometries:

- *“Single-landmark forest”*: Periodic world with only one landmark shape: 10 × 40 mm large cones. Landmarks were placed on the nodes of a triangular grid (**Fig. S1E,G**). Cones were fully hidden by virtual fog at distance greater than 70 mm, gradually emerged from the fog when the fly was closer than 70 mm and came into full contrast at distances smaller than 55 mm, at which point the cones had an angular width of 10.39° at the base and an angular height of 36.03° (**Fig. 1J, Fig. S1G**).
- *“Two-landmark forest”*: Periodic world with two landmark shapes: 10 × 40 mm cones and 8 × 30 mm cylinders. The two types of landmarks were alternatingly placed on the nodes of a Cartesian grid at a distance of 60 mm (**Fig. S1H,I**). The virtual fog started at a distance of 15 mm and reached full coverage at a distance of 45 mm (**Fig. S1I**).

#### Free-walking arena

The free-walking arena design (**Fig. S4A-C, S6B,C**) was inspired by a previous study [50] and built to match lighting, space and landmark-interaction in VR. Specifically, the arena featured a large open space to reduce encounters with walls, and, when needed, small unclimbable objects. The arena consisted of a large circular walking platform (radius 11.4 cm) made from textured matt acrylic (TAP Plastics Inc, San Leandro, CA, USA) surrounded by an acrylic cylinder (inner diameter 22.8 cm, height 17.8 cm) mounted on a laser cut acrylic base. The walking platform could be taken out for cleaning. Coating the arena wall with a siliconizing fluid (Sigmacote from Sigma-Aldrich, St. Louis, MO, USA) prevented wing-clipped flies from walking up the wall. The arena wall was mantled with a white diffusor sheet (V-HHDE-PM06-S01-D01 sample, BrightView Technologies, Durham, NC, USA) and backlit with blue LEDs (470 nm, Super Bright LEDs Inc. St. Louis, MO, USA) mounted on a wire mesh-based scaffold that was fixed onto the arena base. A blue-colored backlight was chosen to match conditions in the VR setup. The light intensity inside the free walking arena measured with a power meter (PM100D with S130C Sensor, Thorlabs Inc, Newton, NJ, USA) for an expected wavelength of 461 nm (peak wavelength in spectrogram) was 0.115 mW/cm^2^ at the arena border and 0.105 mW/cm^2^ in the center.

Four custom-made LED panels mounted below the free walking platform, delivered infrared (IR) backlighting for high-contrast video recordings of the fly’s walking behavior (see below) and red stimulation light (627 nm) for optogenetic stimulation. Each LED panel contained spatially intercalated IR and red (LXM2-PD01-0050, LUXEON Rebel Color, Philips Lumileds Lighting Company, San Jose, CA, USA) LEDs. The four panels were mounted on a water-cooled breadboard (breadboard from Thorlabs Inc, Newton, NJ, USA, liquid cooling system from Koolance, Auburn, WA, USA) and connected to a microcontroller board built around a Teensy 2.0 processor (PJRC.com, LLC Sherwood, OR, USA). Light intensities of the IR and red stimulation LEDs were controlled from a computer using serial communication with the microcontroller. The red LEDs could be individually targeted to control illumination independently in 16 sectors. A cross-shaped light separator reduced light spreading from the illuminated to the non-illuminated LED panels (quadrants). Serial commands to the LED controller were sent through a custom-written Python (version 2.7) program, to ensure temporally precise and repeatable delivery of a light stimulation protocol. Four IR LEDs placed at the corners of the arena base plate were coupled to red illumination in the respective quadrant. These four LEDs served as indicators for the red-light stimulation in videos of the free walking arena (see below, **Fig. S6C**). We measured the relationship between LED controller input to the red LEDs and light intensity at 627 nm (**Fig. S6D,** Thorlabs power meter PM100D with S130C Sensor; in ON quadrant measurements range = 1.3 mW for < 10% and 13 mW above, in OFF quadrants range = 1.3 mW)) in a dark room with two out of four quadrants, i.e. two out of four LED panels, switched on (**Fig. S6B**). With 1 %, 5 % and 10 % current we measured a light intensity of 0.27 mW/cm^2^ (0.01 mW/cm^2^), 1.13 mW/cm^2^ (0.05 mW/cm^2^) and 2.25 mW/cm^2^ (0.11 mW/cm^2^) in the illuminated quadrant, respectively (measurement in the non-illuminated quadrant in brackets).

The flies’ behavior was recorded with BIAS (Basic Image Acquisition Software, version v0p49, IO Rodeo, Pasadena, CA, USA) using a video camera (Flea3 1.3 MP Mono USB3 Vision, Point Grey, Richmond, Canada) placed 120 cm above the walking platform. A lens with 16 mm focal length, low distortion (Edmund Optics Inc, Barrington, NJ, USA, stock #63-245) and a 760 nm IR filter (52 mm, Neewer Technology Ltd., Guangdong, China) were mounted to the camera. Videos were recorded at 12.3 Hz (landmark interaction validation assay) or 20 Hz (quadrant assay) with 1008 × 1008 pixel large images covering an area slightly larger than the diameter of the arena resulting in a spatial resolution of about 40 pixel/cm. The entire rig was placed inside a light-tight black enclosure and temperature and relative humidity were kept at 28-30 °C and 28-32 %.

#### Behavioral assays

##### Fixation assay in VR

All fixation assays were performed at high room temperature (30° C) unless stated otherwise. We performed two types of assays:

- <u>Black stripe fixation</u>: One 10 min trial per fly with the dark-on-bright 20° wide stripe. Through the FlyoVeR GUI visual feedback from translational movements was disabled.
- <u>Comparison of fixation in 1D and 2D scenes with contrast inversion</u>: Each fly was tested in four 10 min trials, presented in random order. In two trials we used the dark-on-bright condition and in the other two the bright-on-dark condition. For each condition, we ran one stripe fixation (1D, translation disabled in FlyoVeR) trial and one trial with the *single-landmark forest* (2D).

##### Validation of landmark interaction in VR

Each fly was exposed to 4 trials (10 min each) in the dark-on-bright *single-landmark forest* VR. In three out of the four trials the landmarks were visible, in one trial they were invisible, i.e., no visual cue was provided as to where the landmark was. The three trials with visible landmarks were always measured in a block with the invisible landmark measured either as the first or last trial.

##### Validation of landmark interaction in freely walking flies

We placed a small 3D-printed black cone, which served as a landmark, in the center of the free walking arena (**Fig. S4A,C**). The cone had the same dimensions as the virtual cones used in the *single-landmark forest* world (10 mm wide, 40 mm high). The surface of the object was polished to reduce surface texture. To prevent flies from climbing and resting on the cone, a coated (SurfaSil, Fisher Scientific, Thermo Fisher Scientific, Waltham, MA, USA) glass cylinder (ø = 15 mm) was placed around the cone. A single wing-clipped fly was introduced to the arena and it was given 1-2 min to explore the space before starting the video recording for a 10 min trial. After each measurement, the fly was removed from the arena and the arena floor was wiped with wet tissue paper.

##### “Virtual sugar” stimulation in freely walking flies

Groups of 10-15 wing-cut female Gr64f-Gal4 > ChrimsonR flies were inserted into free walking arena with ambient blue light (but no objects) and walking behavior was filmed during a 4 min long stimulation protocol. The protocol consisted of 15 pulses (200 ms long and with a light intensity of 1.58 mW/cm^2^), each separated by a 15 s long break. Flies were either wet starved for 24 h in an empty food vial with a humidified filter paper (starved group) or wet starved for 3 h and then transferred back on food for ~1 h (fed group).

##### Quadrant assay in freely walking flies

We used a free walking quadrant assay inspired by [78] to screen stimulation paradigms for their capacity to induce avoidance behavior. HC-GAL4 > ChrimsonR flies were exposed to a stimulation protocol consisting of a 10 s long pre-stimulation period followed by 6 blocks of 30 s of red light stimulation separated by 10 s with no stimulation (**Fig. S6A**). During the 30 s stimulation blocks, red light illumination was restricted to two opposing quadrants and the illuminated quadrants were alternated in consecutive blocks. Per trial responses of all-male or all-female groups of 12-18 wing-clipped flies were measured.

##### Local search in VR

In all local search experiments, we used 24h wet-starved female, wing-cut Gr64f-Gal4 > ChrimsonR flies. For all three experimental groups we used the black-on-bright *single-landmark forest* VR, but landmark visibility and optogenetic stimulation differed:

- <u>Invisible LM + stim</u>: Invisible landmarks and optogenetic stimulation (1.29 mW/cm^2^).
- <u>Visible LM + stim</u>: Visible landmarks and optogenetic stimulation (1.29 mW/cm^2^).
- <u>Only LM:</u> Visible landmarks, but no optogenetic stimulation. For convenience the level of optogenetic stimulation was set to 1%, corresponding to < 0.05 mW/cm^2^, so that flies were not stimulated, but the data could be analyzed in the same way as for the other two groups.

Optogenetic stimulation was triggered whenever the fly crossed the 10 mm visit radius around the center of a landmark. Regardless of what the fly did, optogenetic stimulation lasted for 200 ms. After receiving a stimulation pulse, flies had to leave a 30 mm large zone around the landmark at which the optogenetic stimulation had been triggered, to enable successive stimulation at that site. Whenever flies resided for more than 30 s within 7 mm radial distance from the center of a landmark, they were teleported back to the starting position.

##### “Virtual heat” avoidance in VR

In “virtual heat” avoidance experiments we tested each fly in four 10 min long trials in the *single-landmark forest* world. Three trials with visible landmarks were measured in a block consisting of a *“LM pre”* and *“LM post”* trial, in which the fly was free to explore the purely visual VR, and a *“LM + opt”*, in which each landmark was paired with a *virtual heat* zones (**Fig. 5E**). Either before or after this block, virtual heat avoidance was tested in world with invisible landmarks paired with *virtual heat* zones (*“opt”* trial). Circular *virtual heat* zones had a radius of 40 mm.

##### Visual conditioning in VR

For visual conditioning assay we used the dark-on-bright *two-landmark forest* (**Fig. S1H**). The protocol consisted of three trials, a 10 min *“*Pre*”* trial, a 20 min *“*Training*”* trial and a 10 min *“*Post*”* trial.

<u>Aversive visual conditioning with *virtual heat*:</u> In the aversive visual conditioning experiments, we used male, 5-10d old HC-GAL4 > ChrimsonR flies. In the pre and post trial phases, flies received constant low-level optogenetic stimulation (0.35 mW/cm^2^) independent of their position in the VR. In the training trial *virtual heat* and *virtual cool* zones were introduced by varying optogenetic stimulation levels as a function of the fly’s position in the VR. Both zones had a radial size of 25 mm. In the *cool zones* the stimulation decreased from baseline to 0 mW/cm^2^, while in the *heat zones* the stimulation increased from baseline to 0.81 mW/cm^2^. In all trials, a fly was “teleported” to the start location if it remained within a 7 mm radius around a landmark for more than 30 s.

<u>Appetitive visual conditioning with *virtual sugar*:</u> In appetitive visual conditioning experiments, we used 5-10d old Gr64f > ChrimsonR flies. In pre and post trials, no optogenetic stimulation was provided. In training trials each visit (15 mm radius) to the reward landmark resulted in a 200 ms optogenetic stimulation of 1.29 mW/cm^2^.

### QUANTIFICATION AND STATISTICAL ANALYSIS

#### Preprocessing and data selection criteria

Data from free and tethered walking experiments were treated the same way with exception of movement thresholds and sampling rate. To analyze data from VR trials (logged at 360 Hz), the log file saved by FlyoVeR was parsed with a custom Python (version 2.7) script to extract trial-specific information from the header and the time series data, which was downsampled to 20 Hz using linear interpolation. The locations of objects in the 3D scene used in the respective trial were read from the coordinate file as specified in the log file header. Most of the analysis of walking trajectories in 2D virtual worlds was performed on “collapsed” trajectories, i.e. after pooling trajectory fragments across different locations of the VR that correspond to the same sensory environment, exploiting the periodicity of the virtual world. To obtain collapsed trajectories from trials in the *single-landmark forest*, trajectory fragments within a radial distance of 60 mm each of the periodically placed cone were projected onto a circular area (radius = 60 mm) around the central landmark (**Fig. 1H,I**) preserving the absolute heading direction. Trajectories from trials with the *two-landmark forest* were collapsed in a similar way onto the central square formed by two cylinders and two cones (**Fig. S1H, Movie 1**). Walking trajectories from free walking experiments were extracted from video recordings using Ctrax [86]. The video frame corresponding to the beginning of the optogenetic stimulation protocol in the quadrant assay was determined for each movie using custom macros in Fiji (version 2.0.0-rc-43), by monitoring the brightness of one of the indicator IR LEDs. The trajectory time series from VR and free walking experiments were further analyzed in Python (version 2.7). We classified the behavior of a fly on a per time step basis as “*moving”* if the instantaneous translational velocity exceeded 2 mm/s in VR experiments and 5 mm/s in free walking experiments. On a per trials basis we classify the behavior of a fly as “*walking”* if the fly was moving for more than 20 % of the trial time.

#### Quantification of fixation behavior

To quantify fixation behavior, we employed the following strategy: We first selected all trials in which the fly was *walking* and computed the relative heading angle of the fly with respect to the stripe or the center of the landmark, i.e. the angular location of the stripe or landmark in the fly’s field of view (**Fig. S2C,E**). From this we computed the frequency distribution of relative heading angles for each trial using 20° wide bins (**Fig. S2D,F**). We then attempted to fit the heading distribution with a von Mises distribution (**Fig. S2D**), which is mathematically described as

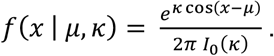

The von Mises distribution is unimodal, i.e. it has a single peak whose location is set by the location parameter μ ∈ (−π,π). The height of the peak is a function of the shape parameter κ. The larger κ, the more mass is centered at the location μ. If κ = 0, the von Mises becomes a uniform circular distribution. If a good fit with a von Mises distribution could be achieved (p > 0.1 with a Kolmogorov-Smirnov test) and the fit had a shape parameter κ > 0.5, we classified a fly’s behavior as “*unimodal fixation*”. Some trials showed a very narrow fixation peak on top of a baseline, which is not well fitted by a von Mises. We therefore also classified a trial as *unimodal fixation*, if the length of the mean angular vector (PVA, population vector average) was larger than 0.5 even if no good fit was achieved. Amongst the heading distributions that did not match the criteria for *unimodal fixation*, several showed two defined peaks (**Fig. S2F**). We therefore added a second step, in which we tried to fit the not yet classified distributions with a bimodal distribution generated by adding two von Mises that share the same shape parameter:

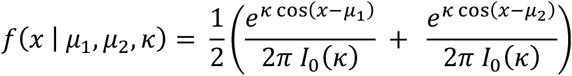

If a good fit with a shape parameter κ > 0.5 could be achieved with this bimodal distribution, the respective trial was classified as “*bimodal fixation*”. All remaining trials were classified as “*unclassified*”. In (**Fig. 2B,F,G**, **S3B,C**) we also made a distinction between trials for which no good fit was achieved (*“No fit”*, p > 0.1), trials that were well fitted with a unimodal or bimodal von Mises distribution, respectively, but did not fulfill the fixation criteria (*“Good fit, no fixation”*, p < 0.1, κ < 0.5 and PVA < 0.5) and those trials that matched fixation criteria (*“Fixation”*)

#### Visit analysis

We quantified “visits” to a landmark as approaches up to a radial distance of a defined “visit radius” or less. We chose a visit radius of 10 mm for local search experiments, the radius at which virtual sugar was delivered, and 15 mm for all other experiments, the radius at all landmarks but the closest ones are hidden. The visit length was defined as the time period between entering and exiting the zone defined by the visit radius.

#### Quantification of local search paths

Our analysis of local search after *virtual sugar* stimulation focused on trajectory fragments (paths) before and after a stimulation event. We only considered paths of a minimal length, 150 mm, that were not truncated by the beginning or the end of the trial or a previous or successive stimulation events, respectively. For the center of mass analysis, paths from all flies were pooled. For the quantification of tortuosity and distance moved, we first averaged paths from a single fly (median) and then across flies. Tortuosity quantified as the path length over the displacement distance within a 2 s sliding window.

#### Statistics

##### Stripe fixation (Fig. 2A,B)

We measured a preference for fixating a stripe in the frontal FOV by testing first whether, for a given group of flies, the fitted location parameter μ was not uniformly distributed using the Rayleigh test. If the Rayleigh test suggested a non-uniform distribution, we computed the circular mean and checked whether it was close to 0° (frontal FOV).

| Genotype | Sex, sample size | Rayleigh test | Circular mean |
| --- | --- | --- | --- |
| DL | Male, n=3 | 2.5906, $p = 0.063$<br>(ns) | - |
| | Female, n = 6 | $z = 5.8045$, $p = 0.0005$ (***) | -0.079 (**-4.529** degrees) |
| WTB hybrid | Male, n = 10 | z = 4.9615, p = 0.0044 (**) | -0.082 rad (-**4.684** degrees) |
|  | Female, n = 14 | z = 4.4454, p = 0.0094 (**) | -0.334 (-**19.137** degrees) |

##### Quadrant assay (Fig. 6D)

Two-sided one-sample t-test (null hypothesis: sample mean = 0.5) over the median residency per experimental repeat within the last 10 seconds of the stimulation block. Statistic performed in Python (version 2.7) using scipy.stats’s ttest_1samp method.

| Stimulation | Male, + Retinal | Female, +Retinal | Male, -Retinal | Female, - Retinal |
| --- | --- | --- | --- | --- |
| 0.27 mW/cm<sup>2</sup> | p = 3.31e-17 (***) | p = 8.39e-10 (***) | p = 0.1853 (ns) | p = 0.5235 (ns) |
| 1.13 mW/cm<sup>2</sup> | p = 4.47e-42 (***) | p = 1.41e-18 (***) | p = 3.18e-10 (***) | p = 0.0472 (*) |
| 2.25 mW/cm<sup>2</sup> | p = 3.29e-37 (***) | p = 1.32e-31 (***) | p = 3.63e-16 (***) | p = 0.0011 (**) |

##### “Virtual heat” avoidance in VR (Fig. S6E)

We used a paired Wilcoxon test (wilcox.test in R, version 3.3.3) to compare the total visit count between two experimental conditions.

We compared (a) data from the two trials with to data from the two trials without optogenetic stimulation and (b) data from the *LM pre* and the *LM post* trial.

| Group | Visit count ~ Optogenetic stim. present / not present | Visit count ~ *LM Pre* / *LM Post* trial |
| --- | --- | --- |
| Group 1: Retinal-fed, no baseline | p = 3.15e-09 (***) | p = 0.3252 (ns) |
| Group 2: Retinal-fed, 0.35 mW/cm<sup>2</sup> baseline | p = 0.0003843 (***) | p = 0.4832 (ns) |
| Group 3: No retinal, no baseline | p = 0.95 (ns) | p = 0.1874 (ns) |

Significance codes: “ns” p > 0.1, “*” p ≤ 0.05, “**” p ≤ 0.01, “***” p ≤ 0.001

### ADDITIONAL RESOURCES

More detailed information about the virtual reality setup and the design of virtual worlds will shortly be provided on www.flyfizz.org.

## References

1. Frisch, K.v. (1967). The dance language and orientation of bees, (Cambridge, Mass.,: Belknap Press of Harvard University Press).

2. Tinbergen, N., and Kruyt, W. (1938). Über die Orientierung des Bienenwolfes (Philanthus triangulum Fabr.). Z Vergl Physiol 25, 292–334.

3. Muller, M., and Wehner, R. (1988). Path integration in desert ants, Cataglyphis fortis. Proc Natl Acad Sci U S A 85, 5287–5290.

4. Wehner, R., and Srinivasan, M.V. (1981). Searching Behavior of Desert Ants, Genus Cataglyphis (Formicidae, Hymenoptera). J Comp Physiol 142, 315–338.

5. Dacke, M., Baird, E., Byrne, M., Scholtz, C.H., and Warrant, E.J. (2013). Dung Beetles Use the Milky Way for Orientation. Current Biology 23, 298–300.

6. Ofstad, T.A., Zuker, C.S., and Reiser, M.B. (2011). Visual place learning in Drosophila melanogaster. Nature 474, 204–207.

7. Collett, T.S., and Collett, M. (2002). Memory use in insect visual navigation. Nat Rev Neurosci 3, 542–552.

8. Wehner, R. (2003). Desert ant navigation: how miniature brains solve complex tasks. J Comp Physiol A Neuroethol Sens Neural Behav Physiol 189, 579–588.

9. Collett, M., Chittka, L., and Collett, T.S. (2013). Spatial Memory in Insect Navigation. Current Biology 23, R789–R800.

10. Dombeck, D.A., and Reiser, M.B. (2012). Real neuroscience in virtual worlds. Current Opinion in Neurobiology 22, 3–10.

11. Stowers, J.R., Hofbauer, M., Bastien, R., Griessner, J., Higgins, P., Farooqui, S., Fischer, R.M., Nowikovsky, K., Haubensak, W., Couzin, I.D., et al. (2017). Virtual reality for freely moving animals. Nat Methods 14, 995–1002.

12. Holscher, C., Schnee, A., Dahmen, H., Setia, L., and Mallot, H.A. (2005). Rats are able to navigate in virtual environments. J Exp Biol 208, 561–569.

13. Dombeck, D.A., Harvey, C.D., Tian, L., Looger, L.L., and Tank, D.W. (2010). Functional imaging of hippocampal place cells at cellular resolution during virtual navigation. Nature Neuroscience 13, 1433–1440.

14. Aronov, D., and Tank, D.W. (2014). Engagement of Neural Circuits Underlying 2D Spatial Navigation in a Rodent Virtual Reality System. Neuron 84, 442–456.

15. Acharya, L., Aghajan, Z.M., Vuong, C., Moore, J.J., and Mehta, M.R. (2016). Causal Influence of Visual Cues on Hippocampal Directional Selectivity. Cell 164, 197–207.

16. Gray, J.R., Pawlowski, V., and Willis, M.A. (2002). A method for recording behavior and multineuronal CNS activity from tethered insects flying in virtual space. Journal of Neuroscience Methods 120, 211–223.

17. Takalo, J., Piironen, A., Honkanen, A., Lempeä, M., Aikio, M., Tuukkanen, T., and Vähäsöyrinki, M. (2012). A fast and flexible panoramic virtual reality system for behavioural and electrophysiological experiments. Scientific Reports 2, 1–9.

18. Kaupert, U., Thurley, K., Frei, K., Bagorda, F., Schatz, A., Tocker, G., Rapoport, S., Derdikman, D., and Winter, Y. (2017). Spatial cognition in a virtual reality home-cage extension for freely moving rodents. J Neurophysiol 117, 1736–1748.

19. Jouary, A., Haudrechy, M., Candelier, R., and Sumbre, G. (2016). A 2D virtual reality system for visual goal-driven navigation in zebrafish larvae. Sci Rep 6, 34015.

20. Wolf, R., and Heisenberg, M. (1990). Visual Control of Straight Flight in Drosophila-Melanogaster. Journal of Comparative Physiology a-Sensory Neural and Behavioral Physiology 167, 269–283.

21. Maimon, G., Straw, A.D., and Dickinson, M.H. (2008). A simple vision-based algorithm for decision making in flying *Drosophila*. Current Biology 18, 464–470.

22. Seelig, J.D., Chiappe, M.E., Lott, G.K., Dutta, A., Osborne, J.E., Reiser, M.B., and Jayaraman, V. (2010). Two-photon calcium imaging from head-fixed *Drosophila* during optomotor walking behavior. Nature methods 7, 535–540.

23. Buchner, E. (1976). Elementary movement detectors in an insect visual system. Biological Cybernetics 24, 85–101.

24. Bahl, A., Ammer, G., Schilling, T., and Borst, A. (2013). Object tracking in motion-blind flies. Nat Neurosci 16, 730–738.

25. van Breugel, F., and Dickinson, M.H. (2012). The visual control of landing and obstacle avoidance in the fruit fly Drosophila melanogaster. J Exp Biol 215, 1783–1798.

26. Saxena, N., Natesan, D., and Sane, S.P. (2018). Odor source localization in complex visual environments by fruit flies. J Exp Biol 221.

27. Kim, I.S., and Dickinson, M.H. (2017). Idiothetic Path Integration in the Fruit Fly Drosophila melanogaster. Curr Biol 27, 2227–2238 e2223.

28. Alvarez-Salvado, E., Licata, A.M., Connor, E.G., McHugh, M.K., King, B.M., Stavropoulos, N., Victor, J.D., Crimaldi, J.P., and Nagel, K.I. (2018). Elementary sensory-motor transformations underlying olfactory navigation in walking fruit-flies. Elife 7.

29. Seelig, J.D., Chiappe, M.E., Lott, G.K., Dutta, A., Osborne, J.E., Reiser, M.B., and Jayaraman, V. (2010). Two-photon calcium imaging from head-fixed Drosophila during optomotor walking behavior. Nat Methods 7, 535–540.

30. Moore, R.J., Taylor, G.J., Paulk, A.C., Pearson, T., van Swinderen, B., and Srinivasan, M.V. (2014). FicTrac: a visual method for tracking spherical motion and generating fictive animal paths. J Neurosci Methods 225, 106–119.

31. Bell, J.S., and Wilson, R.I. (2016). Behavior Reveals Selective Summation and Max Pooling among Olfactory Processing Channels. Neuron 91, 425–438.

32. Schulze, A., Gomez-Marin, A., Rajendran, V.G., Lott, G., Musy, M., Ahammad, P., Deogade, A., Sharpe, J., Riedl, J., Jarriault, D., et al. (2015). Dynamical feature extraction at the sensory periphery guides chemotaxis. Elife 4.

33. Lin, H.-H., Chu, L.-A., Fu, T.-F., Dickson, B.J., and Chiang, A.-S. (2013). Parallel Neural Pathways Mediate CO_2_ Avoidance Responses in *Drosophila*. Science 340, 1338 LP-1341.

34. Klein, M., Afonso, B., Vonner, A.J., Hernandez-Nunez, L., Berck, M., Tabone, C.J., Kane, E.A., Pieribone, V.A., Nitabach, M.N., Cardona, A., et al. (2015). Sensory determinants of behavioral dynamics in *Drosophila* thermotaxis. Proceedings of the National Academy of Sciences 112, E220–E229.

35. Claridge-Chang, A., Roorda, R.D., Vrontou, E., Sjulson, L., Li, H., Hirsh, J., and Miesenböck, G. (2009). Writing Memories with Light-Addressable Reinforcement Circuitry. Cell 139, 405–415.

36. Riemensperger, T., Kittel, R.J., and Fiala, A. (2016). Optogenetics in Drosophila Neuroscience. Methods Mol Biol 1408, 167–175.

37. Aso, Y., and Rubin, G.M. (2016). Dopaminergic neurons write and update memories with cell-type-specific rules. eLife 5, e16135–e16135.

38. Keene, A.C., and Masek, P. (2012). Optogenetic induction of aversive taste memory. Neuroscience 222, 173–180.

39. Stern, U., and Yang, C.-H. (2017). SkinnerTrax: high-throughput behavior-dependent optogenetic stimulation of *Drosophila*. bioRxiv.

40. Corfas, R.A., and Dickinson, M.H. (2018). Diverse food-sensing neurons trigger idiothetic local search in Drosophila. bioRxiv.

41. Brockmann, A., Murata, S., Murashima, N., Boyapati, R.K., Shakeel, M., Prabhu, N.G., Herman, J.J., Basu, P., and Tanimura, T. (2017). Sugar intake elicits a small-scale search behavior in flies and honey bees that involves capabilities found in large-scale navigation. bioRxiv.

42. Murata, S., Brockmann, A., and Tanimura, T. (2017). Pharyngeal stimulation with sugar triggers local searching behavior in Drosophila. J Exp Biol 220, 3231–3237.

43. Ofstad, T.A., Zuker, C.S., and Reiser, M.B. (2011). Visual place learning in Drosophila melanogaster. Nature 474, 204–207.

44. Bahl, A., Ammer, G., Schilling, T., and Borst, A. (2013). Object tracking in motion-blind flies. Nature Neuroscience 16, 730–738.

45. Cohen, J.D., Bolstad, M., and Lee, A.K. (2017). Experience-dependent shaping of hippocampal CA1 intracellular activity in novel and familiar environments. eLife 6.

46. Bülthoff, H., Götz, K.G., and Herre, M. (1982). Recurrent Inversion of Visual Orientation in the Walking Fly, *Drosophila melanogaster*. Journal of Comparative Physiology 148, 471–481.

47. Reichardt, W., and Poggio, T. (1976). Visual control of orientation behaviour in the fly: Part I. A quantitative analysis. Quarterly Reviews of Biophysics 9, 311–311.

48. Reiser, M.B., and Dickinson, M.H. (2008). A modular display system for insect behavioral neuroscience. Journal of Neuroscience Methods 167, 127–139.

49. Heisenberg, M., and Buchner, E. (1977). Role of Retinula Cell-Types in Visual Behavior of Drosophila-Melanogaster. Journal of Comparative Physiology 117, 127–162.

50. Robie, A.A., Straw, A.D., and Dickinson, M.H. (2010). Object preference by walking fruit flies, *Drosophila melanogaster*, is mediated by vision and graviperception. Journal of Experimental Biology 213, 2494–2506.

51. Gallio, M., Ofstad, T.A., Macpherson, L.J., Wang, J.W., and Zuker, C.S. (2011). The coding of temperature in the Drosophila brain. Cell 144, 614–624.

52. Dethier, V.G. (1976). The hungry fly : a physiological study of the behavior associated with feeding, (Cambridge, Mass.: Harvard University Press).

53. Weiss, L.A., Dahanukar, A., Kwon, J.Y., Banerjee, D., and Carlson, J.R. (2011). The molecular and cellular basis of bitter taste in Drosophila. Neuron 69, 258–272.

54. Dahanukar, A., Lei, Y.T., Kwon, J.Y., and Carlson, J.R. (2007). Two Gr genes underlie sugar reception in Drosophila. Neuron 56, 503–516.

55. Fujii, S., Yavuz, A., Slone, J., Jagge, C., Song, X., and Amrein, H. (2015). Drosophila sugar receptors in sweet taste perception, olfaction, and internal nutrient sensing. Curr Biol 25, 621–627.

56. Barbagallo, B., and Garrity, P.A. (2015). Temperature sensation in Drosophila. Curr Opin Neurobiol 34, 8–13.

57. Gallio, M., Ofstad, T.A., Macpherson, L.J., Wang, J.W., and Zuker, C.S. (2011). The Coding of Temperature in the *Drosophila* Brain. Cell 144, 614–624.

58. Ward, S. (1973). Chemotaxis by the nematode Caenorhabditis elegans: identification of attractants and analysis of the response by use of mutants. Proc Natl Acad Sci U S A 70, 817–821.

59. Pierce-Shimomura, J.T., Morse, T.M., and Lockery, S.R. (1999). The fundamental role of pirouettes in Caenorhabditis elegans chemotaxis. J Neurosci 19, 9557–9569.

60. Berg, H.C., and Brown, D.A. (1972). Chemotaxis in Escherichia coli analysed by three-dimensional tracking. Nature 239, 500–504.

61. Dill, M., Wolf, R., and Heisenberg, M. (1993). Visual-Pattern Recognition in Drosophila Involves Retinotopic Matching. Nature 365, 751–753.

62. Aso, Y., Sitaraman, D., Ichinose, T., Kaun, K.R., Vogt, K., Belliart-Guérin, G., Plaçais, P.-Y., Robie, A.a., Yamagata, N., Schnaitmann, C., et al. (2014). Mushroom body output neurons encode valence and guide memory-based action selection in *Drosophila*. eLife 3, e04580–e04580.

63. Nuwal, N., Stock, P., Hiemeyer, J., Schmid, B., Fiala, A., and Buchner, E. (2012). Avoidance of heat and attraction to optogenetically induced sugar sensation as operant behavior in adult Drosophila. Journal of neurogenetics 26, 298–305.

64. Hamada, F.N., Rosenzweig, M., Kang, K., Pulver, S.R., Ghezzi, A., Jegla, T.J., and Garrity, P.a. (2008). An internal thermal sensor controlling temperature preference in *Drosophila*. Nature 454, 217–220.

65. Kitamoto, T. (2001). Conditional modification of behavior in drosophila by targeted expression of a temperature-sensitive shibire allele in defined neurons. Journal of Neurobiology 47, 81–92.

66. Parisky, K.M., Agosto, J., Pulver, S.R., Shang, Y.H., Kuklin, E., Hodge, J.J.L., Kang, K., Liu, X., Garrity, P.A., Rosbash, M., et al. (2008). PDF Cells Are a GABA-Responsive Wake-Promoting Component of the Drosophila Sleep Circuit. Neuron 60, 672–682.

67. Fry, S.N., and Wehner, R. (2005). Look and turn: landmark-based goal navigation in honey bees. J Exp Biol 208, 3945–3955.

68. Barbagallo, B., and Garrity, P.A. (2015). Temperature sensation in Drosophila. Volume 34. pp. 8–13.

69. Zhang, S.W., Lehrer, M., and Srinivasan, M.V. (1999). Honeybee memory: navigation by associative grouping and recall of visual stimuli. Neurobiology of learning and memory 72, 180–201.

70. Judd, S.P.D., and Collett, T.S. (1998). Multiple stored views and landmark guidance in ants. Nature 392, 710–714.

71. Tang, S., Wolf, R., Xu, S., and Heisenberg, M. (2004). Visual pattern recognition in Drosophila is invariant for retinal position. Science 305, 1020–1022.

72. Geva-Sagiv, M., Las, L., Yovel, Y., and Ulanovsky, N. (2015). Spatial cognition in bats and rats: from sensory acquisition to multiscale maps and navigation. Nat Rev Neurosci 16, 94–108.

## ADDITIONAL REFERENCES

73. Azanchi, R., Kaun, K.R., and Heberlein, U. (2013). Competing dopamine neurons drive oviposition choice for ethanol in *Drosophila*. Proceedings of the National Academy of Sciences of the United States of America 110, 21153–21158.

74. Tuthill, John C., Nern, A., Holtz, Stephen L., Rubin, Gerald M., and Reiser, Michael B. (2013). Contributions of the 12 Neuron Classes in the Fly Lamina to Motion Vision. Neuron 79, 128–140.

75. Pfeiffer, B.D., Ngo, T.T.B., Hibbard, K.L., Murphy, C., Jenett, A., Truman, J.W., and Rubin, G.M. (2010). Refinement of tools for targeted gene expression in Drosophila. Genetics 186, 735–755.

76. Schuster, S., Strauss, R., and Gotz, K.G. (2002). Virtual-reality techniques resolve the visual cues used by fruit flies to evaluate object distances. Curr Biol 12, 1591–1594.

77. Triphan, T., Poeck, B., Neuser, K., and Strauss, R. (2010). Visual targeting of motor actions in climbing Drosophila. Curr Biol 20, 663–668.

78. Klapoetke, N.C., Murata, Y., Kim, S.S., Pulver, S.R., Birdsey-Benson, A., Cho, Y.K., Morimoto, T.K., Chuong, A.S., Carpenter, E.J., Tian, Z., et al. (2014). Independent optical excitation of distinct neural populations. Nature Methods 11, 338–346.

79. Isono, K., Tanimura, T., Oda, Y., and Tsukahara, Y. (1988). Dependency on light and vitamin A derivatives of the biogenesis of 3-hydroxyretinal and visual pigment in the compound eyes of Drosophila melanogaster. The Journal of general physiology 92, 587–600.

80. Wang, X., Wang, T., Ni, J.D., von Lintig, J., and Montell, C. (2012). The Drosophila Visual Cycle and De Novo Chromophore Synthesis Depends on rdhB. Journal of Neuroscience 32, 3485–3491.

81. Aso, Y., Hattori, D., Yu, Y., Johnston, R.M., Iyer, N.a., Ngo, T.-T.B., Dionne, H., Abbott, L., Axel, R., Tanimoto, H., et al. (2014). The neuronal architecture of the mushroom body provides a logic for associative learning. eLife 3, 1–47.

82. Land, M.F. (1997). Visual Acuity in Insects. Annual Review of Entomology 42, 147–177.

83. Autrum, H. (1958). Electrophysiological analysis of the visual systems in insects. Experimental Cell Research 14, 14.

84. Cosens, D., and Spatz, H.C. (1978). Flicker Fusion Studies in Lamina and Receptor Region of Drosophila Eye. Journal of Insect Physiology 24, 587-&.

85. Miall, R.C. (1978). The flicker fusion frequencies of six laboratory insects, and the response of the compound eye to mains fluorescent ‘ripple’. Physiol Entomol 3, 99–106.

86. Branson, K., Robie, A.A., Bender, J., Perona, P., and Dickinson, M.H. (2009). High-throughput ethomics in large groups of *Drosophila*. Nature Methods 6, 451–457.

